# KIF2C-induced nuclear condensation concentrates PLK1 and phosphorylated BRCA2 at the kinetochore microtubules in mitosis

**DOI:** 10.1101/2024.04.13.589357

**Authors:** Anastasiia Skobelkina, Manon Julien, Sylvain Jeannin, Simona Miron, Tom Egger, Rady Chaaban, Guillaume Bouvignies, Rania Ghouil, Claire Friel, Didier Busso, François-Xavier Theillet, Romain Le Bars, Aura Carreira, Angelos Constantinou, Jihane Basbous, Sophie Zinn-Justin

**Affiliations:** Institute for Integrative Biology of the Cell (I2BC), CEA, CNRS, Uni Paris-Sud, Uni Paris-Saclay, Gif-sur-Yvette, France; Institut de Génétique Humaine, Université de Montpellier, CNRS, Montpellier, France; Genome Instability and Cancer Predisposition Laboratory, Centro de Biologia Molecular Severo Ochoa (CBMSO), CSIC-UAM, Madrid 28049, Spain; Institut Curie, PSL Research University, CNRS, UMR3348, F-91405, Orsay, France; Laboratoire des Biomolécules, LBM, Département de chimie, École Normale Supérieure, PSL University, Sorbonne Université, CNRS, 24 rue Lhomond, 75005 Paris, France; School of Life Sciences, University of Nottingham, Medical School, QMC, Nottingham, NG7 2UH, United Kingdom; CIGEx, Université Paris Cité et Université Paris-Saclay, INSERM, CEA, Genetic Stability, Stem Cells and Radiation, F-92260 Fontenay-aux-Roses, France

**Keywords:** Mitosis, phosphorylation, condensates, kinetochore, genome stability

## Abstract

During mitosis, the human microtubule depolymerase KIF2C increases the turnover of kinetochore-microtubule attachments. This facilitates the correction of attachment errors. Moreover, BRCA2 phosphorylated at Thr207 by PLK1 (BRCA2-pT207) assembles a complex including PLK1, PP2A and BUBR1 that contributes to the stability of the kinetochore-microtubule attachments. PLK1, together with Aurora B, critically regulate the accurate segregation of chromosomes. Here we demonstrate that KIF2C contains an N-terminal domain that binds directly to several phosphorylated peptides, including BRCA2-pT207. Using an optogenetic platform, we reveal that KIF2C assembles into membrane-less compartments or biomolecular condensates that are located next to microtubules. We provide evidence that condensate assembly depends on the presence of the newly defined N-terminal phospho-binding domain of KIF2C and on the kinase activities of Aurora B and PLK1. Moreover, KIF2C condensates concentrate active PLK1 and colocalize with BRCA2-pT207. We propose that, because of its phospho-dependent binding and oligomerization capacities, KIF2C forms biomolecular condensates that partition PLK1 and locally amplify its kinase activity during mitosis.

**Graphical Abstract:** 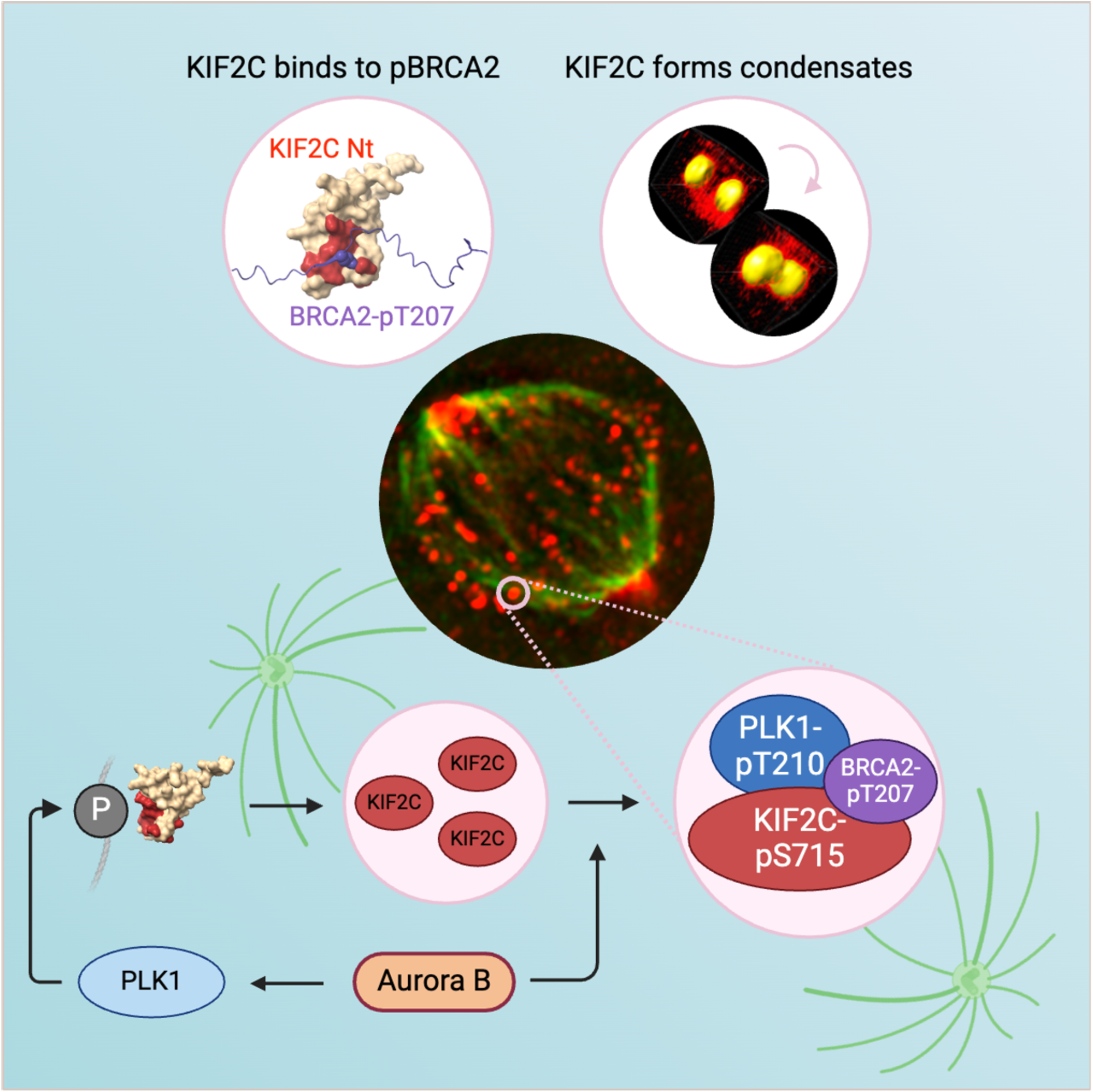

## INTRODUCTION

During mitosis, the proper distribution of chromosomes into daughter cells is a highly controlled process that is essential for genome integrity. Accurate chromosome segregation relies on the bi-oriented attachment of chromosomes to the mitotic spindle. This is ensured through binding of the kinetochores to microtubules oriented towards each spindle pole. During early phases of mitosis, initial capture of kinetochores by microtubules is asynchronous and stochastic, and erroneous attachments are frequent. Persistence of these errors is a common cause of chromosome mis-segregation, associated chromosomal instability and aneuploidy, as observed in solid tumors (Cimini et al., 2001; Siegel and Amon, 2012; Weaver and Cleveland, 2007). Erroneous attachments are converted to bi-oriented attachments to ensure faithful chromosome segregation (Godek et al., 2015). The efficiency of correction depends on the rate of turnover of kinetochore-microtubule attachments: erroneous attachments must be released to enable the formation of new, correct attachments. Cancer cells with chromosomal instability have excessively stable kinetochore-microtubule attachments (Bakhoum et al., 2009b). Strategies that destabilize these attachments promote faithful chromosome segregation in cancer cells by increasing the rate of correction of attachment errors. The microtubule depolymerase KIF2C (alias Mitotic Centromere-Associated Kinesin, MCAK) plays a key role at centromeres/kinetochores by increasing kinetochore-microtubule attachment turnover, thus facilitating error correction prior to anaphase (Hunter et al., 2003; Maney et al., 1998). Overexpression of KIF2C reduces the rate of chromosome mis-segregation (Bakhoum *et al*., 2009b; Kleyman et al., 2014). Inversely, depleting KIF2C is sufficient to promote chromosome segregation defects to levels comparable to those of cancer cells with chromosomal instability (Bakhoum et al., 2009a).

Kinases and phosphatases also contribute to proper chromosome segregation by generating switching phospho-sites that are fine-tuned to correct attachment errors (Saurin, 2018). At the kinetochore-microtubule interface, phosphorylation events are inhibitory to the attachment process, because they electrostatically interfere with microtubule binding (Wimbish and DeLuca, 2020). Phosphorylation of NDC80 by Aurora B thus inhibits the microtubule attachment process. Conversely, Polo-like kinase 1 (PLK1) stabilizes kinetochore-microtubules attachments through phosphorylation of different substrates (Lenart et al., 2007; Li et al., 2010). In particular, it phosphorylates residues S676 and T680 of BUBR1, which enhances recruitment of the B56 subunit of Protein Phosphatase 2A (PP2A-B56), thereby ensuring the correct strength and dynamics of the kinetochore-microtubule attachments (Corno et al., 2023; Elowe et al., 2007; Kruse et al., 2013; Suijkerbuijk et al., 2012). Mitotic regulation by AuroraB-PLK1-PP2A operates only at specific subcellular localization and at certain times, but how this is controlled remains unclear.

Assembly of membrane-less organelles through multivalent intermolecular interactions was recently established as a mechanism that creates high local concentrations, favoring prompt catalysis with precise spatial-temporal control. Increasing evidence suggests that this mechanism contributes to the regulation of microtubule structure and dynamics (Lawrence et al., 2023). Indeed, several proteins interacting with tubulin form condensates *in vitro* and recruit tubulin within these condensates. Tubulin condensation is implicated in spindle apparatus assembly (Jiang et al., 2015), nucleation of acentrosomal and branched microtubules (Hernandez-Vega et al., 2017; King and Petry, 2020) and centrosome maturation (Jiang et al., 2021; Woodruff et al., 2017). Microtubule dynamics is also fine-tuned by Tip-Interacting Proteins. Condensation of these proteins on microtubules, and more specifically at the tip of microtubules, was recently reported *in vitro* and in cells. This process leads to high local concentrations of tubulin and favors microtubule growth (Miesch et al., 2023; Wu et al., 2021).

Recently, we discovered that the tumor suppressor protein BRCA2 is present at the kinetochore in mitosis and contributes to the stability of the kinetochore-microtubule attachments (Ehlen et al., 2020). We and others demonstrated that BRCA2 is phosphorylated by PLK1 (Ehlen *et al*., 2020; Lee et al., 2004; Lin et al., 2003). We identified four PLK1 phospho-sites in the N-terminal region of BRCA2, including S193 and T207 that are conserved from mammals to fishes (Ehlen *et al*., 2020; Julien et al., 2021; Julien et al., 2020b). Phosphorylation of T207 triggers a direct interaction between BRCA2 (specifically BRCA2-pT207) and PLK1, and the assembly of a complex between BRCA2, PLK1, BUBR1 and PP2A. This complex is essential for the stability of the kinetochore-microtubule interactions and the proper alignment of chromosomes (Ehlen *et al*., 2020). Reduced binding of phosphorylated BRCA2 to PLK1, as observed in *BRCA2* breast cancer variants S206C and T207A, results in erroneous chromosome segregation and aneuploidy. In this study, we identified a direct, PLK1-dependent, interaction between KIF2C and BRCA2-pT207, which is mediated by a previously unrecognized KIF2C phospho-peptide binding domain. In mitotic cells, KIF2C assembles into biomolecular condensates that are located adjacent to microtubules and kinetochores. These condensates are enriched in activated PLK1 as well as in phosphorylated BRCA2. We propose that KIF2C condensates favor timely phosphorylation by PLK1 of proteins located at the kinetochore-microtubule interface in mitosis.

## RESULTS

### BRCA2 phosphorylated at T207 interacts with the microtubule depolymerases MCAK/KIF2C and KIF2A

We searched for phospho-dependent partners of BRCA2 in mitosis. We focused on partners of BRCA2 phosphorylated by PLK1 at two highly conserved sites: S193 and T207. Therefore, we produced and purified two batches of recombinant Avi-tagged BRCA2_167-260_, and phosphorylated one of these batches with PLK1. We verified using Nuclear Magnetic Resonance (NMR) that S193 and T207 were totally phosphorylated (Figure 1A). The less conserved T219 and T226 were also phosphorylated, as reported (Ehlen *et al*., 2020). Following our recently published protocol (Bouguechtouli et al., 2024), we then (i) biotinylated the two batches (phosphorylated and non-phosphorylated) using BirA, (ii) attached the resulting peptides on streptavidin-coated magnetic beads and (iii) incubated the beads in mitotic human cell extracts. Proteins interacting with either non-phosphorylated or phosphorylated BRCA2_167-260_ were identified and quantified using mass spectrometry. The resulting plot showed that PLK1 was the most enriched protein on the phospho-peptide coated beads (Figure 1B, left panel). We performed a similar experiment using BRCA2_167-260_ mutated at T207. By comparing the lists of proteins binding to the phospho-peptides BRCA2_167-260_ WT and T207A, we observed that PLK1 interacts with BRCA2_167-260_ only in the presence of phosphorylated T207 (pT207) (Figure 1B, right panel). This is fully consistent with our previously published crystal structure of the complex between the Polo-Box Domain of PLK1 and a peptide fragment of BRCA2 phosphorylated at T207 (Ehlen *et al*., 2020). We identified two other proteins from the Volcano plot analysis: the microtubule depolymerases MCAK/KIF2C and KIF2A (Figure 1B). Interaction with these proteins also involved pT207: peptides corresponding to KIF2C and KIF2A were almost twice more enriched on beads with phosphorylated BRCA2_167-260_ WT as compared to T207A. All BRCA2 binding partners, *i.e.* PLK1, KIF2C and KIF2A, were not detected when beads were incubated in asynchronous human cell extracts (Suppl. Fig. 1A).

**Figure 1.**
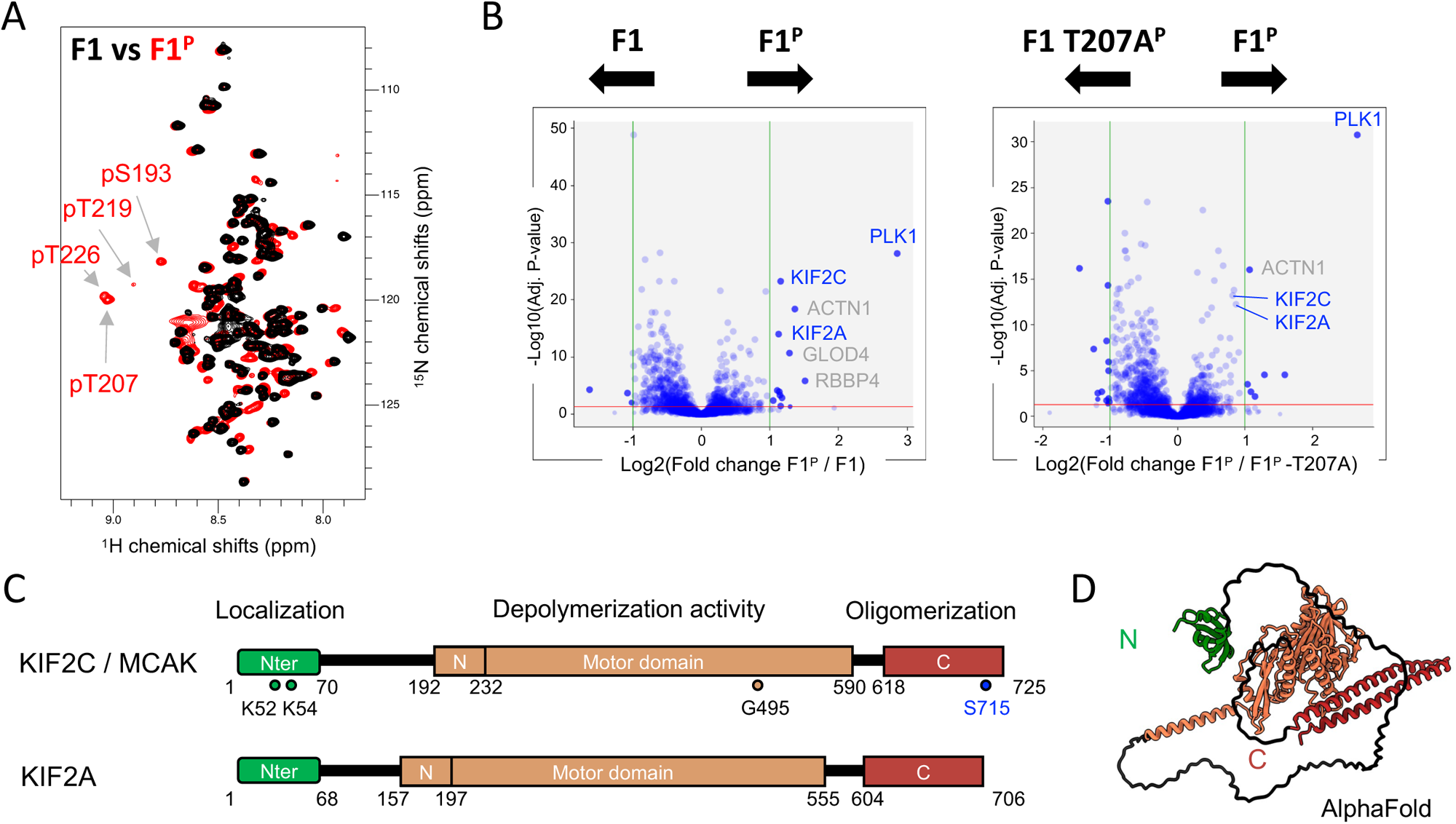
BRCA2 phosphorylated by the mitotic kinase PLK1 interacts with the microtubule depolymerases KIFC and KIF2A. **(A)** NMR characterization of BRCA2_167-270_ before (F1; black) and after phosphorylation by PLK1 (F1^P^; red). Superposition of the 2D NMR ^1^H-^15^N SO-FAST HMQC spectra of F1 and F1^P^, where each peak represents the chemical environment of a BRCA2 residue, identified phosphorylated residues; peaks corresponding to the four phosphorylated residues are labeled. **(B)** Volcano plots showing proteins from mitotic HeLa cell extracts that were identified by mass spectrometry as binding to (left) BRCA2_167-270_ phosphorylated by PLK1 (F1^P^) versus non phosphorylated BRCA2_167-270_ (F1), and (right) BRCA2_167-270_ phosphorylated by PLK1 (F1^P^) versus phosphorylated BRCA2_167-270_ variant containing the T207A mutation (F1 T207A^P^). **(C)** Domain organization of KIF2C and KIF2A, with residues of KIF2C mutated in this study indicated by colored circles: K52 and K54 in the N-terminal domain, G495 in the catalytic site, and S715 involved in KIF2C dimerization. **(D)** AlphaFold model of KIF2C, colored as in (C).

### Phosphorylated BRCA2_167-260_ binds directly to the N-terminal domains of KIF2C and KIF2A

The sequences of the two depolymerases KIF2C and KIF2A are 53% identical (Suppl. Fig. 1B). As predicted by AlphaFold, KIF2C and KIF2A exhibit an uncharacterized N-terminal folded domain, a large disordered region, a neck and motor domain that is responsible for the catalytic activity of the enzymes (Maney et al., 2001; Ogawa et al., 2004), and an α-helical C-terminal region that controls the oligomerization of KIF2C (Zhang et al., 2011) (Figure 1C, D; Suppl. Fig. 1B). The motor domain of KIF2C interacts with tubulin (Wang et al., 2017) and dimerizes upon binding to a C-terminal peptide of KIF2C centered on S715 (Suppl. Fig. 1C; (Talapatra et al., 2015)). We tested direct binding of phosphorylated BRCA2_167-260_ to the different domains of KIF2C. Therefore, we superimposed 2D NMR ^1^H ^15^N spectra of this construct recorded in the absence and presence of KIF2C domains. We could not detect any interaction between phosphorylated BRCA2_167-260_ and the neck and motor domain of KIF2C in our conditions (Suppl. Fig. 2A), whereas we observed a clear interaction between phosphorylated BRCA2_167-260_ and the N-terminal domain of KIF2C (KIF2C Nt) (Figure 2A). NMR ^1^H-^15^N peaks corresponding to the BRCA2 region T203-R212 showed a strong (larger than 60%) decrease in intensity upon addition of KIF2C Nt, demonstrating that this region interacts directly with KIF2C Nt. We also showed that phosphorylated BRCA2_167-260_ interacts with the N-terminal domain of KIF2A (KIF2A Nt) through the same BRCA2 region (Figure 2B). To further identify BRCA2 residues whose chemical environment is most affected upon binding and confirm the central role of phosphorylated T207, we performed an NMR ^15^N CEST experiment. We used a BRCA2_48-218_ construct that contains only two PLK1 phosphorylated sites (S193 and T207), hence avoiding overlap between NMR peaks of phosphorylated residues. By superimposing 2D NMR ^1^H ^15^N spectra of this construct recorded in the absence and presence of KIF2C Nt, we observed that several NMR peaks, including that of S205, pT207, L209 and V211, disappeared upon addition of KIF2C Nt. Furthermore, the ^15^N CEST experiment revealed that the ^15^N frequency difference between BRCA2 free and bound states is particularly large for these 4 residues, demonstrating that they form the core of the binding motif. Finally, we did not detect any interaction between non-phosphorylated BRCA2_167-260_ and the N-terminal domain of either KIF2C or KIF2A (Suppl. Figs. 2B-C). Altogether, we concluded that BRCA2 interacts with KIF2C and KIF2A N-terminal domains through a S_205_S(pT)VLIV_211_ motif centered on phosphorylated T207.

**Figure 2.**
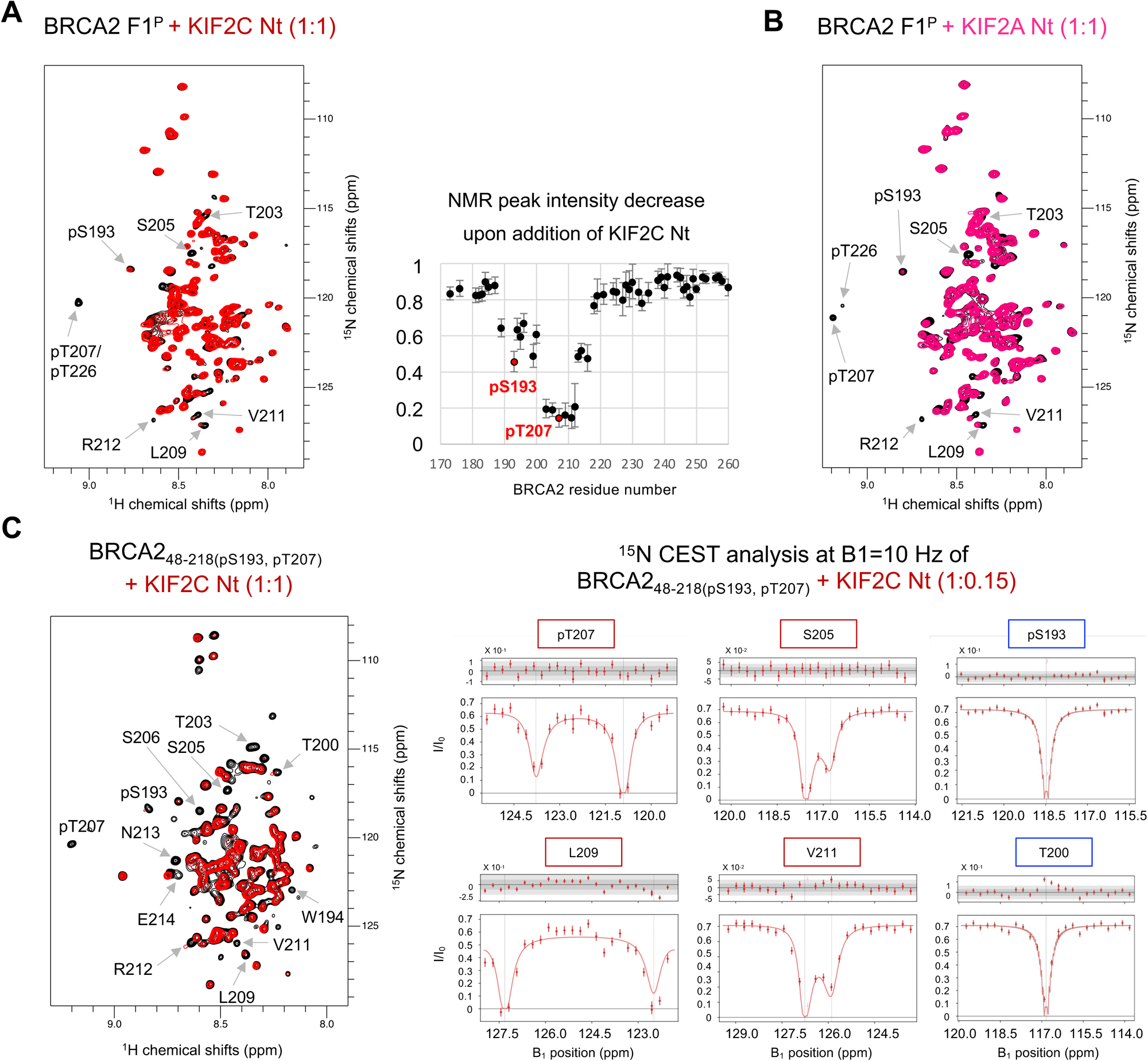
BRCA2 pT207 binds to the N-terminal domains of KIF2C and KIF2A. **(A)** NMR analysis of the interaction between BRCA2_167-260_ phosphorylated by PLK1 (F1^P^) and the N-terminal domain of KIF2C (KIF2C Nt). 2D NMR ^1^H-^15^N SO-FAST HMQC spectra of F1^P^, where each peak represents the chemical environment of a BRCA2 residue, were recorded before (black) and after (red) addition of KIF2C Nt (ratio 1:1). Superimposition of these spectra revealed that, upon binding, several peaks disappeared, while other peaks shifted (marked with arrows). The associated graph displays the peak intensity decrease upon addition of KIF2C Nt as a function of the BRCA2 residue number. **(B)** NMR analysis of the interaction between BRCA2_167-260_ phosphorylated by PLK1 (F1^P^) and the N-terminal domain of KIF2A (KIF2A Nt). Here again, the 2D NMR ^1^H-^15^N SO-FAST HMQC spectra of phosphorylated Avi-tagged BRCA2_167-260_ recorded before (black) and after (magenta) addition of the N-terminal domain of KIF2A (KIF2A Nt; ratio 1:1) were superimposed. The peaks affected by the interaction are labeled. **(C)** NMR characterization of the KIF2C-bound state of phosphorylated BRCA2_48-218_. On the left, 2D NMR ^1^H-^15^N SO-FAST HMQC spectra acquired on phosphorylated BRCA2_48-218_ in the presence and absence of the N-terminal domain of KIF2C (ratio 1:1) were superimposed; several peaks, including those corresponding to S205, pT207, L209 and V11, disappeared after addition of KIF2C. On the right, ^15^N CEST analysis performed on phosphorylated BRCA2_48-218_ in the presence of the KIF2C N-terminal domain (ratio 1:0.15); the major and minor dips observed for residues S205, pT207, L209 and V211 indicate the ^15^N chemical shifts in the free and bound states, respectively.

### The N-terminal domains of KIF2C and KIF2A exhibit a conserved and positively charged cavity that is responsible for phospho-peptide binding

We further characterized the interfaces between phosphorylated BRCA2 and these kinesins. KIF2C and KIF2A N-terminal domains share 43% of identity and 68% of similarity (Suppl. Fig. 1B). KIF2C Nt is monomeric, as shown by Size-Exclusion Chromatography coupled to Multi-Angle Light Scattering (SEC-MALS) (Figure 3A). We measured by Isothermal Titration Calorimetry (ITC) that BRCA2_194-214(pT207)_, a BRCA2 peptide centered on pT207, binds to KIF2C Nt with a K_d_ of 0.48 ± 0.04 µM (Figure 3B; Suppl. Fig. 2D-F; Table 1). It binds to KIF2A Nt with a similar K_d_ of 0.60 ± 0.10 µM. Moreover, we confirmed by ITC that this binding is phospho-dependent (Suppl. Fig. 2E). To identify KIF2C and KIF2A residues involved in BRCA2 binding, we assigned the 2D NMR ^1^H-^15^N HSQC peaks of both N-terminal domains (Suppl. Figs. 3A-D), and we measured the changes in chemical shifts (chemical shift perturbations or CSP) of these peaks upon addition of BRCA2_194-214(pT207)_ (Figure 3C, D; Suppl. Figs. 3E, F). We identified 11 KIF2C residues and 12 KIF2A residues with CSP values larger than 0.2 ppm. Half of these residues are located in the C-terminal β-strand of the β-barrel structure predicted by AlphaFold (Figure 3C, D; Suppl. Figs. 3G, H). KIF2C Nt is significantly less conserved in evolution (from mammals to fishes) than KIF2A Nt (Suppl. Figs. 3A-B). In the case of KIF2C, we could identify a conserved site at the surface of the N-terminal domain (Figure 3E). The BRCA2 binding site as defined by NMR strongly overlaps with the KIF2C Nt conserved site, which is also positively charged (Figure 3E). We performed a similar chemical shift analysis upon addition of non-phosphorylated BRCA2_194-214_ and did not detect any binding between KIF2C or KIF2A Nt and this peptide (Suppl. Fig. 3I). Using AlphaFold Multimer, we further calculated 15 models of the complexes between KIF2C or KIF2A Nt and BRCA2_194-214_. 10 and 14 models have medium interface scores (0.7 > ipTM > 0.5), respectively. All these models showed a β-sheet formed by the KIF2C β-strand A50-D57 (or KIF2A β-strand D48-D55) and the BRCA2 peptide (Figure 3F). In this β-sheet, KIF2C K52 and K54 (or KIF2A K50 and K52) are located in front of phosphorylated T207. These lysines were consistently identified as part of the BRCA2 binding site by NMR (Figures 3C, D). We also produced a mutant construct of the KIF2C N-terminal domain in which K52 and K54 were both mutated into glutamic acid. NMR analysis showed that mutations do not affect the overall structure of the domain, but completely disrupt the binding to BRCA2_194-214(pT207)_ (Suppl. Fig. 3J, K).

**Figure 3.**
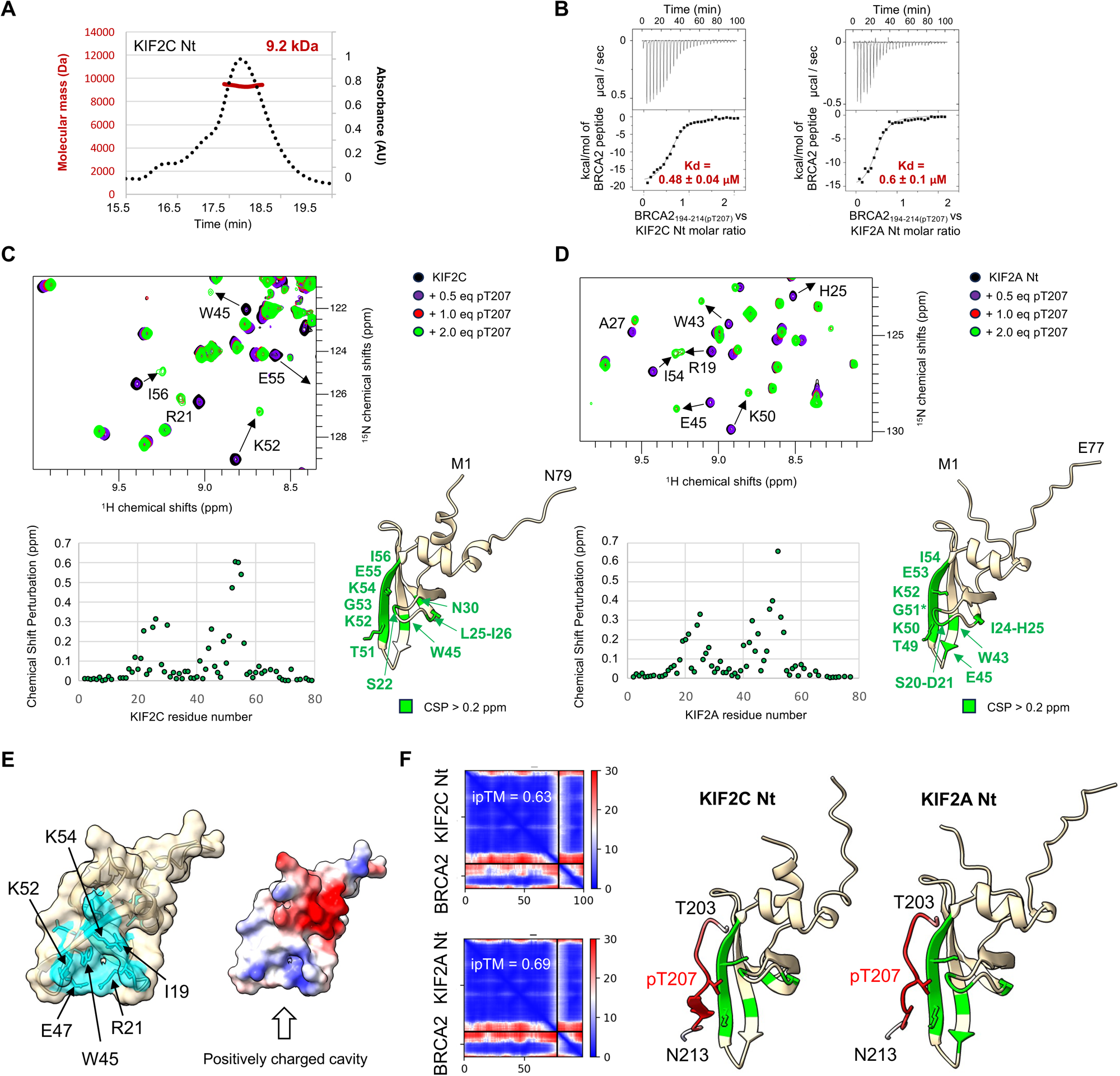
A conserved and positively charged cavity in the N-terminal domains of KIF2C and KIF2A is responsible for phosphorylated BRCA2 peptide binding. **(A)** Determination of the oligomeric state of the N-terminal domain of KIF2C (KIF2C Nt) by SEC-MALS. **(B)** Affinity and stoichiometry of the interaction between KIF2C/KIF2A Nt and BRCA2_194-214(pT207)_, as measured by ITC. **(C)** Identification of the KIF2C Nt residues interacting with BRCA2_194-214(pT207)_ by NMR. Superposition of the 2D NMR ^1^H-^15^N SO-FAST HMQC spectra of KIF2C Nt recorded upon addition of increasing concentrations of BRCA2_194-214(pT207)_ identified the peaks that shifted upon binding. These shifts were quantified by calculating the NMR chemical shift perturbation (CSP) as a function of the KIF2C residue number. The spatial distribution of the residues with CSP values larger than 0.2 ppm is displayed on an AlphaFold model of the N-terminal domain of KIF2C. **(D)** Identification of the KIF2A Nt residues interacting with BRCA2_194-214(pT207)_ by NMR. Here again, the 2D NMR ^1^H-^15^N SO-FAST HMQC spectra of KIF2A Nt recorded upon addition of increasing concentrations of BRCA2_194-214(pT207)_ were superimposed. The NMR chemical shift perturbations (CSP) were plotted as a function of KIF2A residue number. The spatial distribution of the residues with CSP values larger than 0.2 ppm is displayed on an AlphaFold model of the N-terminal domain of KIF2A. G51 is marked by a star because it is not reported in the graph (poor estimation of its CSP). **(E)** Representation of the surface properties of the N-terminal domain of KIF2C. On the left, surface conservation from human to fish is shown, with 100% identical residues in the sequence alignment of Suppl. Figure 3A in cyan. On the right, the electrostatic potential at the surface of the KIF2C N-terminal domain is presented (blue - positively charged - to red - negatively charged). **(H)** Models of the complexes between KIF2C/KIF2A Nt (colored as in panels C and D) and BRCA2_194-214_, as calculated by AlphaFold. Only BRCA2 residues from T203 to N213 are displayed for clarity. These residues are colored as a function of their pLDDT score (red to white: high to low precision on residue position). The corresponding heat maps (representing the error in the relative positioning of the residues of the complex) are shown on the left, and the interface scores (ipTM, from 0 – low confidence – to 1 – high confidence) are indicated.

**Table 1:** Summary of the ITC data obtained by adding synthetic phosphorylated BRCA2/Sgo2 peptides to KIF2C/KIF2A recombinant proteins.

| | Kd (μM) | N | $\Delta H$ (kcal/mol) | $\Delta G$ (kcal/mol) | $-T\Delta S$ (kcal/mol) |
| --- | --- | --- | --- | --- | --- |
| BRCA2<sub>194-214</sub>(pT207) vs KIF2C Nt expt 1 | $0.48 \pm 0.04$ | 0.62 | -19.1 | -19.8 | -0.7 |
| BRCA2<sub>194-214</sub>(pT207) vs KIF2C Nt expt 2 | $1.30 \pm 0.80$ | 0.77 | -12.1 | -12.4 | -0.3 |
| BRCA2<sub>194-214</sub>(pT207) vs KIF2A Nt expt 1 | $0.60 \pm 0.10$ | 0.51 | -15.7 | -16.2 | -0.5 |
| BRCA2<sub>194-214</sub>(pT207) vs KIF2A Nt expt 2 | $0.80 \pm 0.10$ | 0.47 | -18.1 | -18.8 | -0.7 |
| Sgo2<sub>531-544</sub>(pT537) vs KIF2C Nt expt 1 | $0.18 \pm 0.02$ | 0.60 | -9.8 | -9.8 | 0.0 |
| Sgo2<sub>531-544</sub>(pT537) vs KIF2C Nt expt 2 | $0.33 \pm 0.06$ | 0.83 | -8.8 | -8.9 | 0.0 |
| Sgo2<sub>531-544</sub>(pT537) vs KIF2A Nt expt 1 | $0.1 \pm 0.01$ | 0.50 | -16.7 | -17.2 | -0.5 |
| Sgo2<sub>531-544</sub>(pT537) vs KIF2A Nt expt 2 | $0.23 \pm 0.07$ | 0.85 | -10.6 | -10.7 | -0.1 |
| Sgo2<sub>614-627</sub>(pT620) vs KIF2C Nt expt 1 | $0.75 \pm 0.08$ | 0.78 | -21.7 | -22.6 | -0.9 |
| Sgo2<sub>614-627</sub>(pT620) vs KIF2C Nt expt 2 | $0.87 \pm 0.07$ | 0.57 | -11.9 | -12.2 | -0.3 |
| Sgo2<sub>614-627</sub>(pT620) vs KIF2A Nt expt 1 | $0.37 \pm 0.04$ | 0.92 | -14.9 | -15.3 | -0.4 |
| Sgo2<sub>614-627</sub>(pT620) vs KIF2A Nt expt 2 | $0.33 \pm 0.03$ | 0.87 | -15.2 | -15.6 | -0.4 |

We decided to explore the phospho-binding properties of KIF2C and KIF2A N-terminal domains. It was known from the literature that phosphorylation of Shugoshin 2 (Sgo2) by Aurora B at T537 and T620 promotes binding to KIF2C and further localization of KIF2C to centromeres (Figure 4A; (Tanno et al., 2010)). We identified some sequence similarities between BRCA2_194-214(pT207)_ and these Sgo2 phospho-sites (Figure 4B). Therefore, we tested whether the N-terminal domains of KIF2C and KIF2A bind to phosphorylated Sgo2. Using ITC, we observed that both domains interact with Sgo2_531-544(pT537)_ and Sgo2_614-627(pT620)_ with micromolar affinities (Figure 4C; Suppl. Fig. 4A). Using NMR, we showed that these interactions are mediated by the conserved and positively charged surface of KIF2C and KIF2A (Figure 4D; Suppl. Figs. 4B-C). No interaction was observed with the non-phosphorylated Sgo2 peptides by NMR (Suppl. Fig. 4D). Also, no interaction was observed between KIF2C Nt K52E/K54E and Sgo2_531-544(pT537)_ or Sgo2_614-627(pT620)_ (Suppl. Fig. 4E). Finally, AlphaFold Multimer predicted that the Sgo2 peptides form an intermolecular β-sheet with KIF2C and KIF2A, similar to that predicted for BRCA2_(pT207)_ (Figure 4E). Altogether, we concluded that KIF2C and KIF2A Nt interact with phosphorylated peptides sharing the motif pT-X-Φ−Φ, Φ being a hydrophobic residue (Figure 4B). All the tested peptides also contain an arginine, which forms a salt bridge with a negatively charged residue in KIF2C and KIF2A.

**Figure 4.**
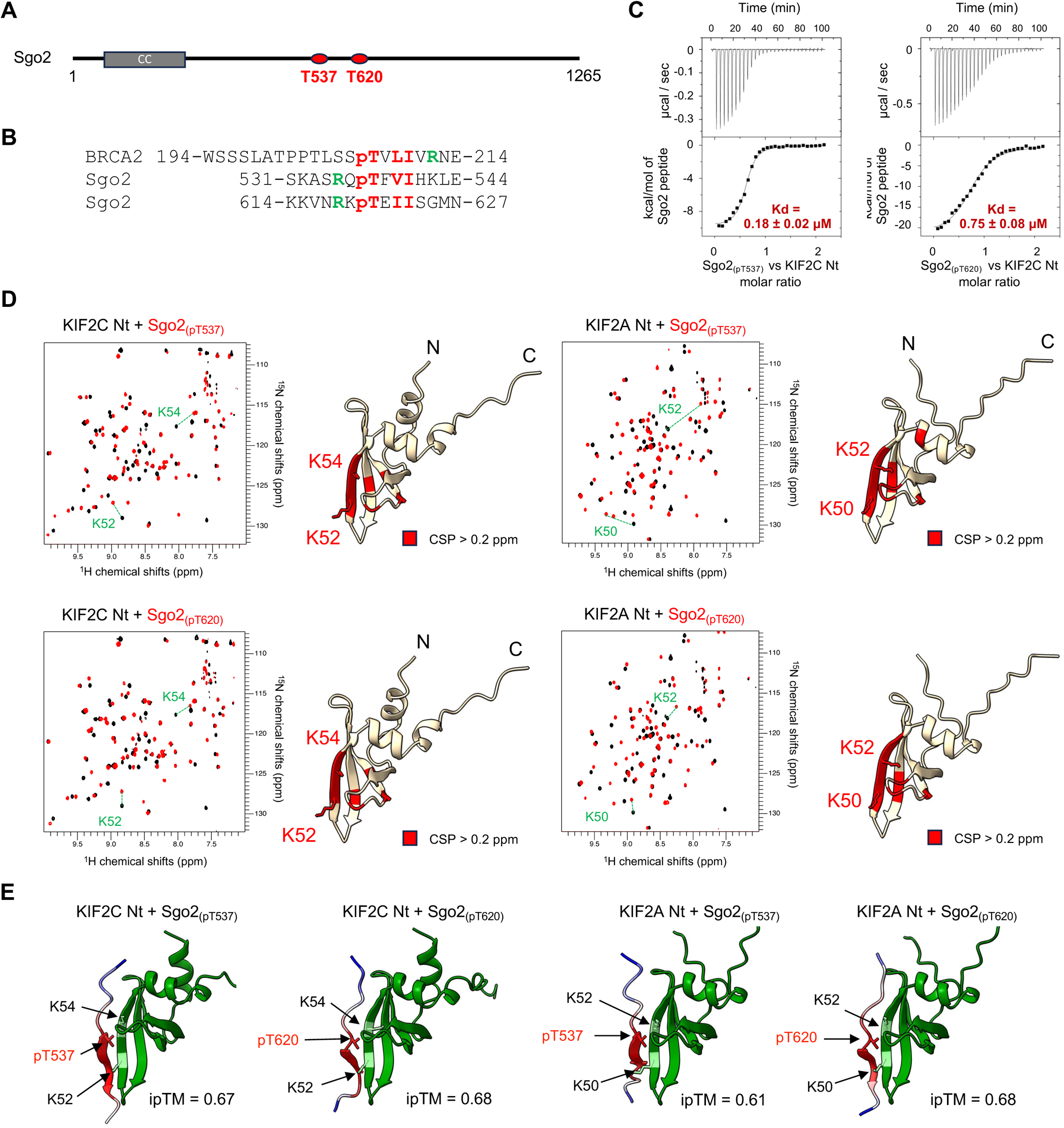
The N-terminal domains of KIF2C and KIF2A bind to a phosphorylated motif also found in Sgo2. **(A)** Scheme of the sequence of human Sgo2, that is responsible for KIF2C recruitment at the centromeres. Phosphosites at T537 and T620 were reported in (Tanno *et al*., 2010). **(B)** Alignment of the sequences of the three putative KIF2C / KIF2A binding peptides. Residues similar in the 3 sequences are in red, those proposed to make a salt bridge with KIF2C or KIF2A by AlphaFold are in green (R212 in BRCA2 with E44 in KIF2C and E42 in KIF2A; R535 and R618 in Sgo2 with E55 in KIF2C and D55 in KIF2A). **(C)** Affinity and stoichiometry of the interaction between KIF2C Nt and Sgo2_(p537)_ or Sgo2_(pT620)_, as measured by ITC. Data obtained with KIF2A are displayed in Suppl. Figure 4A. **(D)** Identification of the KIF2C Nt residues interacting with Sgo2_(pT537)_ or Sgo2_(pT620)_ by NMR. Superposition of the 2D NMR ^1^H-^15^N SO-FAST HMQC spectra of KIF2C/KIF2A Nt recorded before (black) and after addition of Sgo2_(pT537)_ or Sgo2_(pT620)_ (red; ratio 1:2) and mapping of the associated chemical shift perturbations (CSP) onto an AlphaFold model of KIF2C/KIF2A Nt. CSP values are plotted as a function of the residue number in Suppl. Figure 4B. As an example, the 4 spectra of the titration of KIF2C with Sgo2_(pT620)_ are superimposed in Suppl. Figure 4C. **(E)** Models of the complexes between KIF2C/KIF2A Nt (green; K52 and K54 in light green) and Sgo2_(p537)_ or Sgo2_(pT620)_, as calculated by AlphaFold. Sgo2 residues are colored as a function of their pLDDT score, as calculated by AlphaFold (red to blue: high to low precision for residue position).

### The N-terminal domain of KIF2C mediates the formation of KIF2C condensates in cells

Endogenous KIF2C forms foci in mitotic cells, and some of these foci colocalize with kinetochores / centromeres (Wordeman and Mitchison, 1995). Having shown that KIF2C contains a phosphorylated peptide binding domain, and knowing that KIF2C is phosphorylated by various mitotic kinases such as CDK1, Aurora A, Aurora B and PLK1 (Andrews et al., 2004; Lan et al., 2004; Sanhaji et al., 2010; Zhang *et al*., 2011; Zhang et al., 2008), we asked whether KIF2C assembles into condensates through phospho-dependent intermolecular interactions in cells. To investigate this, we fused KIF2C-mCherry to the light-responsive oligomerization domain *Arabidopsis* cryptochrome 2 (Cry2) (Figure 5A) (Alghoul et al., 2021; Bugaj et al., 2013; Frattini et al., 2021). Such optogenetic system makes it possible to control the nucleation of biomolecular condensates in space and time, and thus to characterize the molecular events associated with condensation. We first induced the expression of our KIF2C construct (named Opto-KIF2C) with doxycycline in Flp-In HEK293 cells. We observed that, in the absence of blue light, Opto-KIF2C forms foci in the nuclei of these cells (Figure 5B, line 1; Suppl. Fig. 5A); moreover, after addition of nocodazole, the foci seem brighter and/or larger (Figure 5B, line 2). Upon exposure of the cells (both without and with nocodazole) to an array of blue light LEDs during 5 min of light-dark cycles (4 s light followed by 10 s dark), the number of KIF2C foci per nucleus increased significantly (Suppl. Fig. 5B), and the foci seemed even brighter and/or larger (Figure 5B, lines 3-4). We quantified their fluorescence intensity: the mean fluorescence intensity of the foci increased with illumination (Suppl. Fig. 5C), while the intensity of the free protein in the nucleus decreased (Suppl. Fig. 5D), suggesting that the free protein is recruited to the foci upon illumination. We measured the dimensions of the foci in both non-mitotic and mitotic cells using Structured Illumination Microscopy (SIM) after exposure to 5 min of 488 nm light, and found an averaged foci area of about 0.1 μm^2^ (Figure 5C; Suppl. Fig. 5E). We also demonstrated that addition of nocodazole significantly increased the area of the optogenetic KIF2C condensates (Figure 5C; Suppl. Fig. 5E).

**Figure 5.**
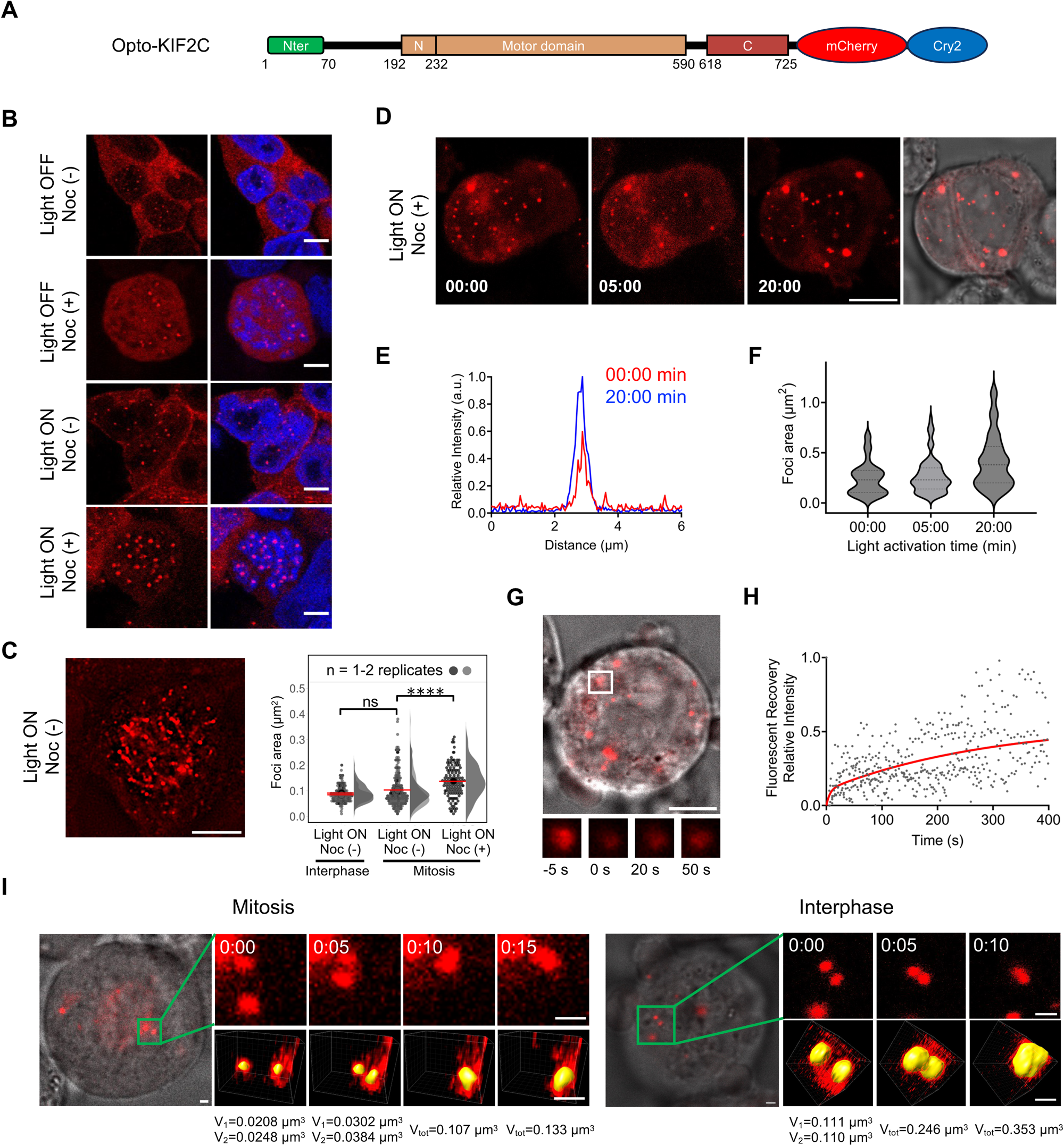
KIF2C forms nuclear membrane-less condensates in cells. **(A)** KIF2C construct used for optogenetic experiments. Opto-KIF2C was used to express KIF2C-mCherry-Cry2 and study KIF2C assembly mechanisms (Alghoul *et al*., 2021). **(B)** Representative fluorescence images obtained after expression of Opto-KIF2C induced by doxycycline in different conditions. In lines 3 and 4, cells were illuminated with blue light (Light ON) during 5 min of light-dark cycles (4 s light followed by 10 s dark). In lines 2 and 4, nocodazole was added to the cell culture (Noc (+)). DNA is stained with Hoechst 33258. Scale bars: 5 µm. **(C)** Structured Illumination Microscopy (SIM) image obtained with the Opto-KIF2C construct on a mitotic cell in light ON and Noc (−) conditions, and quantification of the foci area from a set of SIM images acquired after a 5 min illumination. The p-value calculated under light ON and Noc (−) conditions between mitotic and non-mitotic cells is 0.4270, whereas the p-value calculated under light ON conditions for mitotic cells in the absence (Noc (−)) and presence (Noc (+)) of nocodazole is lower than 0.0001. More representative images are shown in Suppl. Figure 5E. Scale bars: 5 µm. **(D)** Time-lapse microscopy images recorded after addition of nocodazole (Noc (+)) and activation of Opto-KIF2C foci (light ON) in living cells. Time 0: after a first cycle of 15 s light and 30 s dark. Times 5 and 20 (in min): after further illumination by cycles of 8 s light and 30 s dark. Image on the right: merged fluorescence and DIC images acquired after 20 min. Scale bar: 10 µm. **(E)** Line scan showing the increase in size and fluorescence intensity of a selected condensate after 20 min of illumination. **(F)** Violin plot quantification of the condensate area during blue light activation. **(G)** Merged fluorescence and DIC images of a representative cell analyzed by FRAP. Scale bar: 5 μm. **(H)** Quantification of the FRAP data from 6 different experiments. The red curve represents the mean recovery in those experiments. **(I)** Time-lapse 3D microscopy images of activated opto-KIF2C condensates (light ON) in mitosis (left) and in interphase (right). See also Movie S1. Scale bars: 1 µm. Time in sec.

To better understand the nature of these condensates, we performed fluorescence recovery after photobleaching (FRAP) measurements just after illumination. To properly detect fluorescence recovery, we needed to increase the size of the foci. For this purpose, we added nocodazole, illuminated the living cells for one cycle of 15 s light followed by 30 s dark, and then exposed a plane of about 0.5 µm thick showing a large number of foci with light-dark cycles of 8 s light followed by 30 s dark. We observed that, upon blue light exposure, the size and fluorescence signal of KIF2C nuclear foci are enhanced in this plane (Figures 5D-E). Several foci reached an area of about 1 µm^2^ after 20 min of 8 s / 30 s cycling (Figure 5F). In these conditions, we could perform a FRAP analysis of their dynamics by illuminating half of one of the foci and measuring the fluorescence recovery of the protein within it (Figures 5G-H). Although we observed variability between points, mostly due to the different sizes of the foci and the difficulty to bleach part of such highly mobile objects, we found that fluorescence in these foci recovered at 50 % in average with a time constant of about 300 s, which could correspond to a membrane-less condensate recovery time scale in cells. We also observed several fusion events in interphase and mitotic cells (without nocodazole) in the different microscopy movies. To characterize these fusion events, we performed 3D imaging, which allowed us to distinguish between a fusion and a superposition of condensates. We observed that the condensates resulting from fusion exhibited a volume equal to the sum of the volumes of the two initial condensates and regained a circular shape (Figure 5I; Movie S1). Moreover, optogenetic KIF2C condensates fused and continued to grow from surrounding molecules. The rapidly evolving morphology of these condensates confirmed that they are membrane-less cellular sub-compartments.

In order to characterize the molecular mechanisms of KIF2C condensate assembly, we designed five mutants: (i) a variant deleted from its N-terminal domain (Opto-KIF2C_ΔNt_), (ii) K52E/K54E, which lost its phospho-binding capacity (Opto-KIF2C_K52E/K54E_), (iii) S715A, which lost a characterized mitotic PLK1 phosphorylation site (Opto-KIF2C_S715A_), (iv) S715E (Opto-KIF2C_S715E_), which mimics phosphorylation of S715 that disrupts a KIF2C intermolecular interaction (Suppl. Fig. 1C), and (v) G495A, which is catalytically inactive (Opto-KIF2C_G495A_) (Figure 6A). We tested whether these mutants formed condensates in mitotic cells upon blue light exposure: Opto-KIF2C_ΔNt_ and Opto-KIF2C_K52E/K54E_ completely lost their ability to form foci, whereas the other mutants could still assemble into condensates to different degrees (Figures 6B-C; Suppl. Fig. 6A). In the presence of nocodazole, S715E strongly reduced the number of KIF2C condensates, whereas G495A had only a mild impact, and S715A had no impact at all. The same results were obtained in the absence of nocodazole, both for mitotic and non-mitotic cells, except that G495A significantly decreased the number of Opto-KIF2C foci when microtubules were present in the cells (Figure 6B-C; Suppl. Figs 6A-C). In parallel, we tested the impact of PLK1 activity on KIF2C foci formation. We observed that supplementation with a PLK1 inhibitor significantly reduced the number of KIF2C nuclear foci (Figures 6D-E). Even the (small) number of foci formed by KIF2C S715E decreased upon addition of a PLK1 inhibitor (Suppl. Figs. 6D-E). Altogether, our results demonstrated that the lysines of the KIF2C N-terminal domain involved in phospho-peptide binding are key to the assembly of KIF2C condensates in cells. Phosphorylation by PLK1 is also necessary to establish these condensates. Mutation S715E, reported to disrupt a KIF2C intermolecular interaction, decreases KIF2C condensate formation.

**Figure 6.**
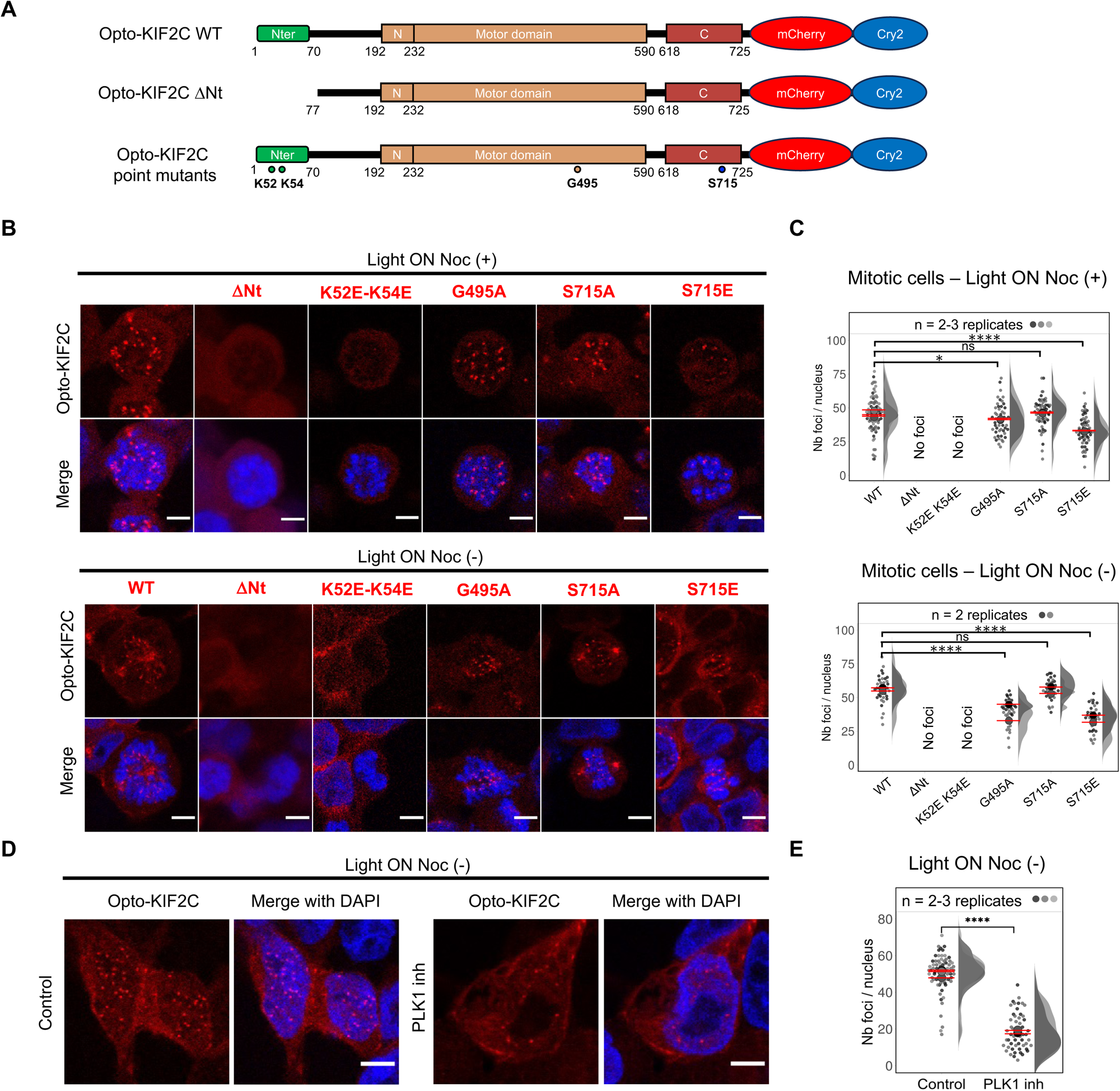
KIF2C assembly into condensates depends on its phosphorylated peptide binding capacity as well as the activity of the mitotic kinase PLK1. **(A)** Constructs used for optogenetic experiments, containing KIF2C either WT, deleted from its N-terminal domain, or with point mutations: K52E/K54E, G495A, S715A, S715E. **(B)** Representative fluorescence images obtained after induction by doxycycline of the expression of Opto-KIF2C mutants and illumination of the cells with blue light (light ON). In the two upper lines, nocodazole was added to the cells (Noc (+)) to observe mostly mitotic cells. In the two lower lines, mitotic cells were analyzed in Noc (−) conditions. Scale bars: 5 µm. **(C)** Quantification of the number of foci per cell as measured on images recorded as in panel (B). In Noc (+) conditions, the p-value corresponding to S715E is lower than 0.0001, whereas those of G495A and S715A are 0.0394 and 0.5616, respectively. In Noc (−) conditions, p-values corresponding to G495A and S715E are lower than 0.0001, whereas that of S715A is 0.7856. **(D)** Impact of PLK1 inhibition on Opto-KIF2C WT foci formation (light ON Noc (−)). **(E)** Quantification of the number of foci per cell as measured on the images recorded as in panel (D). The p-value is lower than 0.0001.

### KIF2C condensates contain BRCA2 phosphorylated at T207 but are next to microtubules and kinetochores

To further describe KIF2C condensates, we searched for KIF2C partners recruited within these condensates. As we revealed that, *in vitro*, KIF2C binds to BRCA2-pT207, we first verified that KIF2C binds to BRCA2-pT207 in mitotic cells. We performed GFP-trap pulldowns using DLD1 BRCA2 deficient cells stably expressing GFP-MBP-BRCA2. We observed that GFP-MBP-BRCA2 co-immunoprecipitated with endogenous KIF2C in mitosis (Figure 7A). Moreover, KIF2C and BRCA2-pT207 co-localized at the kinetochores in metaphase chromosome spreads (Figures 7B-C). Using immunofluorescence microscopy, we found that Opto-KIF2C and BRCA2-pT207 co-localized in condensates assembled upon blue light exposure in Flp-In HEK293 cells (Figures 7D-E). Furthermore, all Opto-KIF2C variants forming condensates co-localized with BRCA2-pT207 (Suppl. Figures 7A-B), except Opto-KIF2C S715A (Figures 7D-E; Suppl. Figure 7B). It was previously reported that KIF2C S715 is phosphorylated by PLK1 in nocodazole conditions (Shao et al., 2015). We concluded that phosphorylation of S715 might be important for BRCA2-pT207 to be localized within Opto-KIF2C condensates. We further tested the presence of tubulin in the condensates, because it is the best characterized KIF2C partner: it directly interacts with the KIF2C motor domain (Wang *et al*., 2017). Unexpectedly, we found that Opto-KIF2C condensates did not co-localize with tubulin (Figure 7F). Opto-KIF2C condensates were positioned adjacent to microtubules, as confirmed using SIM microscopy, and after addition of nocodazole, they excluded tubulin (Figure 7G; Movie S2; Suppl. Figure 7C). Condensates of Opto-KIF2C mutated at S715 were similarly found next to microtubules (Suppl. Figure 7D). We concluded that Opto-KIF2C can interact with tubulin only at the periphery of the condensates. Finally, we tested whether Opto-KIF2C condensates were positioned at the kinetochores in cells. Here again, we found using SIM microscopy that the condensates did not co-localize with the centromere marker CREST, but were positioned next to CREST foci (Figure 7H). Altogether, we established that, when we stimulate the assembly of the Opto-KIF2C condensates using our optogenetic set up, these condensates form adjacent to both microtubules and kinetochores.

**Figure 7.**
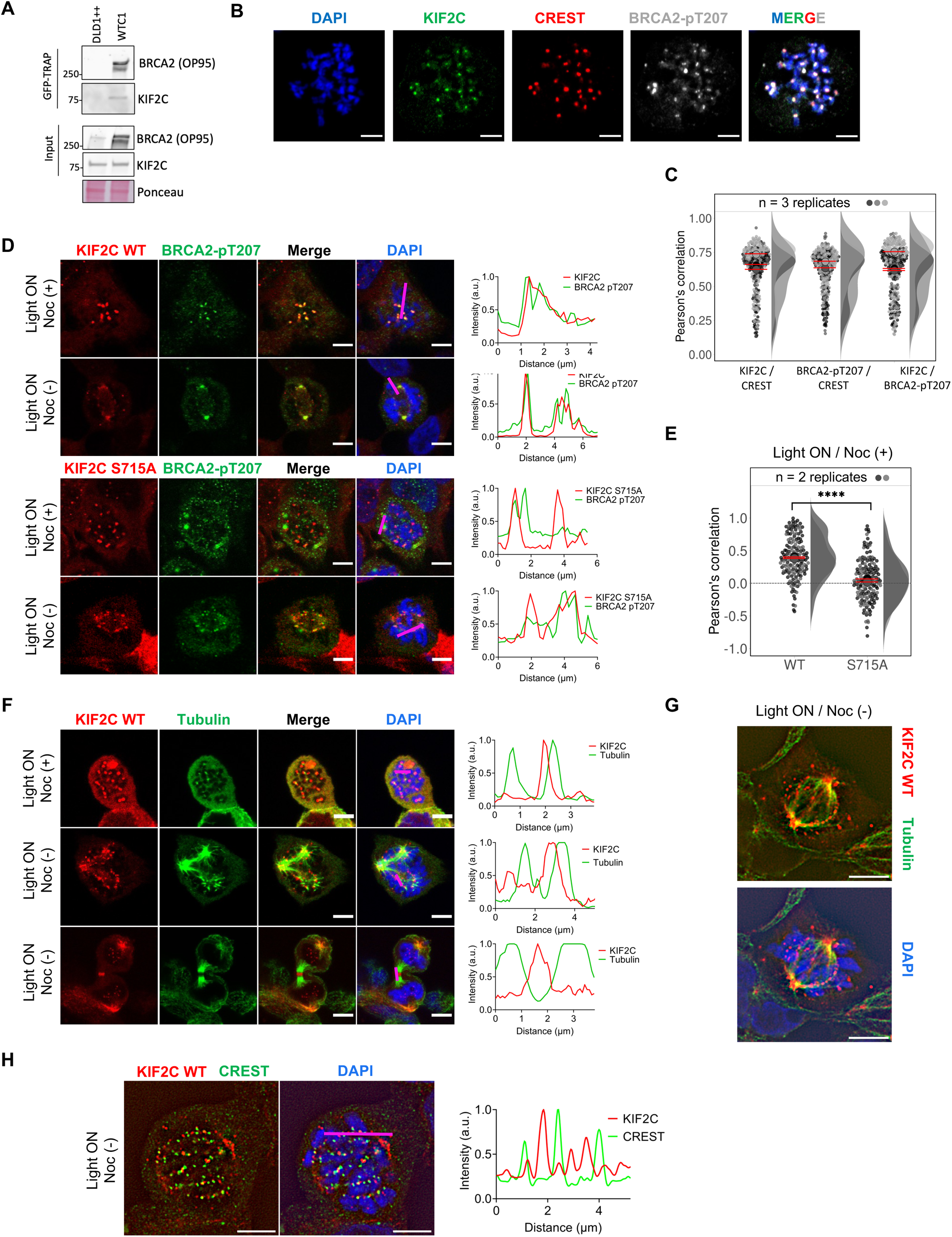
KIF2C condensates concentrate phosphorylated BRCA2 but are found next to microtubules and kinetochores in cells. **(A)** GFP-trap pull-down experiments performed in mitotic DLD1 BRCA2 deficient cells stably expressing GFP-MBP-BRCA2, showing that GFP-MBP-BRCA2 forms a complex with endogeneous KIF2C. The parental DLD1 cell line (DLD1++) was used as a negative control. **(B)** Co-localization of KIF2C, BRCA2-pT207 and the centromere marker CREST observed on metaphase chromosome spreads from DLD1+/+ cells. **(C)** Quantification of the co-localization of KIF2C, BRCA2-pT207, and CREST in metaphase chromosome spreads, analyzed as shown in panel (B). **(D)** Representative immunofluorescence images of Opto-KIF2C WT or S715A (red channel), BRCA2-pT207 (green channel) and DNA (blue channel) obtained in light ON and either Noc (+) or Noc (−) conditions. Line scans on the right show co-localization. **(E)** Quantification, using Pearson’s correlation coefficient, of the co-localization between the indicated proteins (Opto-KIF2C WT or S715A) and BRCA2-pT207, as observed in (D) after addition of nocodazole (Noc (+)). Median shown in red. The p-value is lower than 0.0001. **(F)** Representative immunofluorescence images of Opto-KIF2C WT (red channel), tubulin (green channel) and DNA (blue channel) in light ON conditions (in the presence (Noc (+)) or absence (Noc (−)) of nocodazole). Line scans on the right show that KIF2C foci are next to tubulin. **(G)** SIM images obtained with Opto-KIF2C (red), tubulin (green) and DAPI (blue) in light ON and Noc (−) conditions. See also Movie S2. **(H)** SIM images obtained with Opto-KIF2C (red), CREST (green) and DAPI (blue) in light ON and Noc (−) conditions. The line scan on the right shows that KIF2C foci are next to centromeres, as marked by CREST. Scale bars: 5 µm.

### Functional consequences of KIF2C phosphorylation

To investigate the function of KIF2C condensates, we searched for proteins that localized in the vicinity of KIF2C only after condensate assembly. To do so, we tagged the N-terminus of our Opto-KIF2C construct with TurboID, an optimized biotin ligase that executes promiscuous biotinylation of nearby proteins within minutes (Figure 8A; (Branon et al., 2018)). We induced TurboID-Opto-KIF2C condensation using blue light in the presence of biotin in the cell culture medium and then purified biotinylated proteins with streptavidin-coated beads. We found that, upon 15 min of blue light exposure, PLK1, PLK1-pT210 and tubulin were specifically enriched within KIF2C condensates (Figure 8B; Suppl. Figure 8A). Addition of a PLK1 inhibitor altered the proximity of these proteins to KIF2C foci (Figure 8B; Suppl. Figure 8A). Using immunofluorescence microscopy, we found that Opto-KIF2C and PLK1, as well as activated PLK1 (PLK1-pT210), form similar patterns in all conditions (Figure 8C; Suppl. Figure 8B). When quantifying their co-localizations in nocodazole conditions, we obtained a particularly high Pearson’s correlation coefficient for Opto-KIF2C and PLK1-pT210 (Suppl. Figure 8C-D). We did not observe any strong impact of mutating KIF2C S715 on the co-localization of Opto-KIF2C with PLK1 and PLK1-pT210. Also, following the increase in the number of Opto-KIF2C foci upon blue light exposure (Suppl. Fig. 5B), the number of PLK1 and PLK1-pT210 foci was significantly higher upon illumination (Figure 8D).

**Figure 8.**
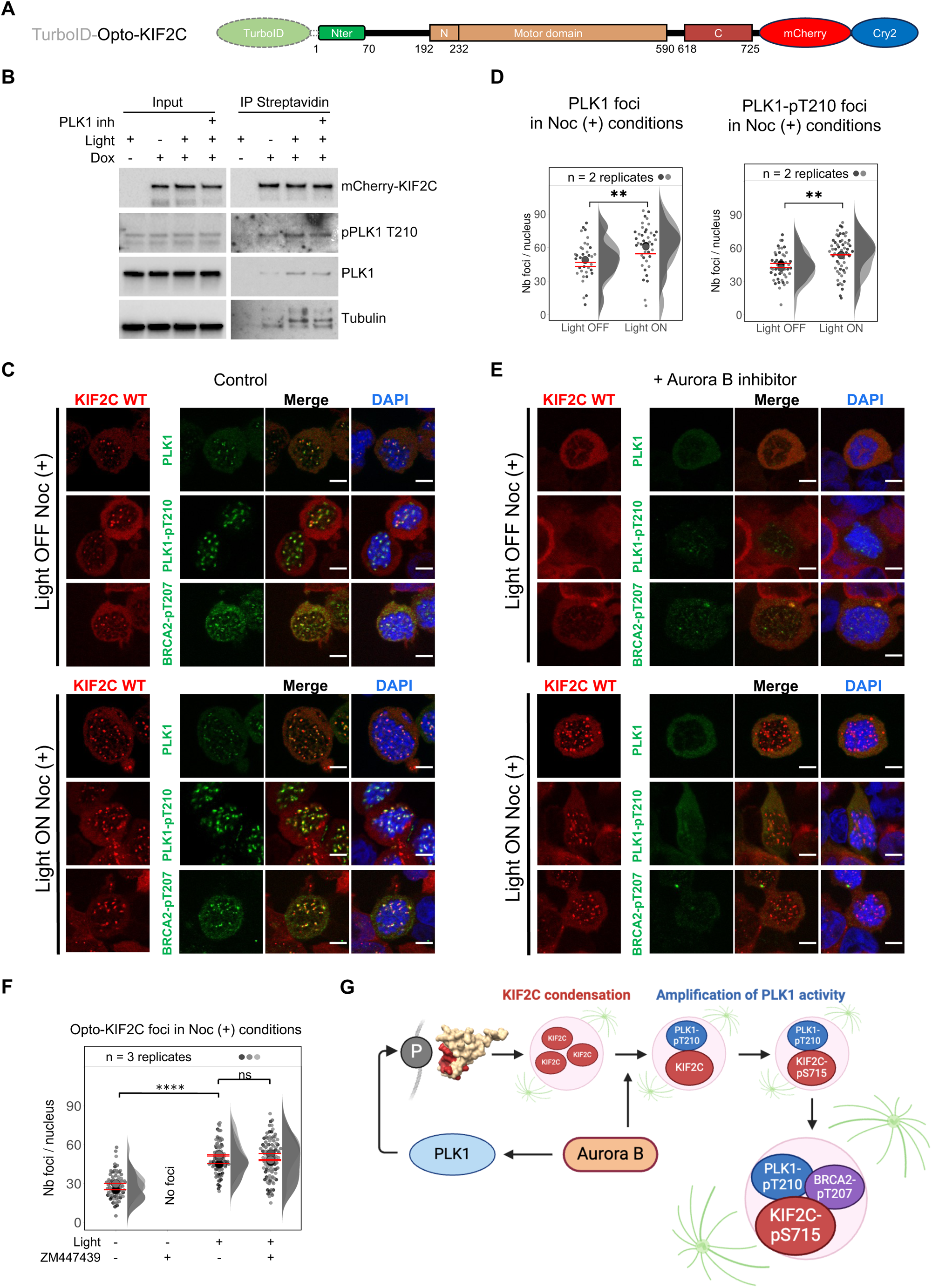
Activated PLK1 is enriched within KIF2C condensates. **(A)** KIF2C construct used for TurboID experiments. **(B)** Identification of TurboID-Opto-KIF2C partners by Western Blot. Partners of TurboID-Opto-KIF2C were biotinylated in different doxycycline and light conditions and purified using streptavidin-coated beads. PLK1, PLK1-pT210 and tubulin were identified as enriched in KIF2C condensates. **(C)** Representative immunofluorescence images of Opto-KIF2C WT (red channel), PLK1 or PLK1-pT210 (green channel) and DNA (blue channel) in nocodazole (Noc (+)) and either light OFF or light ON conditions. Scale bars: 5 µm. **(D)** Quantification of the number of PLK1 and PLK1-pT210 foci per cell as measured on the images recorded as in panel (C). P-values: 0.0015 and 0.0012 for PLK1 and PLK1-pT210, respectively. **(E)** Representative immunofluorescence images obtained as in (C) but upon addition of the Aurora B inhibitor ZM447439. Scale bars: 5 µm. **(F)** Quantification of the number of Opto-KIF2C foci observed in panels (C) and (D). The p-value between the distributions of foci numbers under Light OFF and Light ON conditions is lower than 0.0001, whereas the p-value between Light ON and Light ON + ZM447439 conditions is 0.6951. (**G)** Model for PLK1 regulation in KIF2C condensates. KIF2C condensation is Aurora-B and PLK1-dependent, and through an Aurora B-dependent mechanism, triggers local concentration of PLK1, as well as further phosphorylation of proteins involved in the control of kinetochore-microtubule attachment, including KIF2C-S715 and BRCA2-T207. As KIF2C condensates are next to microtubules, the KIF2C depolymerase activity, enhanced by phosphorylation of S715, might be spatially limited to the periphery of the condensates.

Finally, as Aurora B is required for PLK1 activation at kinetochores (Carmena et al., 2012) and regulates KIF2C at the mitotic centromere (Andrews *et al*., 2004), we checked if the Aurora B inhibitor ZM447439 affected KIF2C condensation and co-localization with PLK1. We observed that, in mitotic cells, the addition of the Aurora B inhibitor completely abolished the formation of KIF2C condensates formed without light (Figure 8E), whereas it either did not change (Noc (+); Figures 8D-F) or mildly decrease (Noc (−); Suppl. Figures 8B, E) the number of optogenetic KIF2C condensates formed upon blue light exposure. Thus, the Aurora B activity is essential for KIF2C condensation, and artificially favoring KIF2C condensation bypasses the requirement for Aurora B activity. We also observed that, upon addition of ZM447439, PLK1 was less abundant in the vicinity of KIF2C condensates in both light OFF and ON conditions (Suppl. Figure 8F). Using immunofluorescence, we were unable to detect PLK1 and PLK1-pT210 within Opto-KIF2C condensates in mitotic cells, both in the absence (Suppl. Fig. 8B) and in the presence of nocodazole (Figure 8E). Similarly, we could not detect BRCA2-pT207 in KIF2C condensates upon addition of the Aurora B inhibitor (Figure 8E). We concluded that Aurora B not only facilitates KIF2C condensate formation, but also promotes the accumulation of PLK1, PLK1-pT210 and BRCA2-pT207 in these condensates (Figure 8G).

## DISCUSSION

During mitosis, KIF2C accumulates in foci, some of which colocalize with kinetochores / centromeres (Wordeman and Mitchison, 1995). The activity of KIF2C at the kinetochores is regulated by Aurora B and PLK1 through multiple feedback loops (Joukov and De Nicolo, 2018). The kinase cascades converge on activation of PLK1, which phosphorylates KIF2C on S715, thus promoting its microtubule depolymerase activity; this event is essential to ensure the timely correction of aberrant kinetochore-microtubule attachments and a proper chromosome segregation (Maney *et al*., 1998) (Hunter *et al*., 2003; Shao *et al*., 2015). Moreover, PLK1 phosphorylates BRCA2 T207, which triggers direct binding between PLK1 and BRCA2, as well as the assembly of a complex between BRCA2, PLK1, PP2A and BUBR1 phosphorylated by PLK1 on T680 (Ehlen *et al*., 2020). This complex plays a critical role in stabilizing kinetochore-microtubule attachments and promoting accurate chromosome alignment. Here, we revealed that KIF2C is able to form membrane-less organelles (Figure 5). It also directly binds to BRCA2 phosphorylated at T207 through its N-terminal domain (Figures 1-4). This domain is essential for KIF2C condensate formation (Figure 6B-C). KIF2C condensation strongly depends on PLK1 activity (Figure 6D-E), and KIF2C condensates concentrate PLK1 (Figure 8B-C). This suggests that KIF2C condensation amplifies PLK1 activity at the kinetochores.

### KIF2C contains a newly described phosphorylated peptide binding domain

We previously revealed that, in mitosis, BRCA2-pT207 interacts with the Polo-Box domain of PLK1, and that the resulting BRCA2-PLK1 complex controls the proper attachment between kinetochores and microtubules (Ehlen *et al*., 2020). In particular, we showed that the BRCA2 variant T207A that has a reduced affinity for PLK1 impairs kinetochore-microtubule attachment stability. While searching for new partners of phosphorylated BRCA2 in mitosis, we found out that the microtubule depolymerases KIF2C and KIF2A exhibit an N-terminal domain binding to BRCA2 phosphorylated at T207 (Figure 2). AlphaFold predicts that the phospho-peptide binding domain of KIF2C has the same barrel-like fold as TUDOR/PWWP domains, which bind to methylated peptides (Gayatri and Bedford, 2014; Maurer-Stroh et al., 2003; Selenko et al., 2001). All these domains interact with their targets through the same cavity. However, whereas this cavity is negatively charged in the case of TUDOR/PWWP domains, it is positively charged in the case of KIF2C and KIF2A. The N-terminal domains of KIF2C and KIF2A bind phosphorylated motifs from different proteins, including BRCA2 and Sgo2 (Figures 3-4). AlphaFold models of the complexes systematically show extended peptides from BRCA2 or Sgo2 forming an antiparallel β-sheet with KIF2C (or KIF2A), the side chain of T207 being close to those of K52 and K54 of KIF2C (K50 and K52 of KIF2A). These models, even if calculated with non-phosphorylated peptides, suggest that phosphorylated T207 interacts through a salt-bridge with one of these lysines. Our NMR mapping of the interfaces is consistent with these models. Mutating K52 and K54 completely abolished KIF2C condensate assembly, indicating that binding of the N-terminal domain of KIF2C to either its own phosphorylated motifs (Ritter et al., 2015) or the phosphorylated motifs present in its partners, as for example BRCA2 or Sgo2, is essential for condensate formation.

### KIF2C forms membrane-less organelles through interactions involving its N-terminal domain, its C-terminal residue S715 and PLK1-dependent phosphorylated sites

Endogenous KIF2C forms foci in mitosis (Wordeman and Mitchison, 1995). In our conditions, KIF2C condensates were observed in cell nuclei even without activation of the optogenetic system (Figure 5B). Upon exposure to 488 nm light, we found that condensates fused and continued to grow from surrounding molecules (Figure 5I; Movie S1). In nocodazole conditions, we were able to quantify the impact of illumination with blue light on the fluorescence intensity and size of the foci. We found that light exposure triggered further recruitment of surrounding KIF2C into the condensates that got slightly brighter (Suppl. Figs. 5B, D) and larger (Figure 5F). We measured that their fluorescence recovery time was about 300 s, as also reported for other membrane-less organelles in cells (Bodmer et al., 2023). We then searched for the molecular determinants of KIF2C condensation in mitosis. We analyzed the condensation mechanisms after adding nocodazole to increase the number of mitotic cells. However, we systematically checked whether similar observations could be found in mitotic cells under Noc (−) conditions, as microtubule depolymerization could affect KIF2C-related processes. First, condensation was efficient when KIF2C S715 was replaced by an alanine and thus could not be phosphorylated, but it was less efficient when this residue was replaced by the phosphomimetic glutamic acid (Figures 6A-B; Suppl. Figs. 6B-C). Structural studies have revealed that S715 phosphorylation disrupts the interaction between two KIF2C motor domains and its C-terminal S715-containing region (Talapatra *et al*., 2015). Our data suggests that the interaction between KIF2C motor and C-terminal regions promotes KIF2C condensate assembly. Additionally, PLK1 activity contributes to KIF2C condensation, as the inhibition of PLK1 significantly decreases the assembly of both KIF2C WT and KIF2C S715E foci (Figures 6C-D; Suppl. Figs. 6D-E). Hence, PLK1 phosphorylates either KIF2C at sites different from S715 (Ritter *et al*., 2015; Sanhaji et al., 2014) or KIF2C partners, such as BRCA2, thereby favoring KIF2C condensation.

### KIF2C condensates contain PLK1 and its phosphorylated targets

Our biotin-proximity labelling and immunofluorescence approaches showed that KIF2C condensation triggers the recruitment of PLK1 (Figures 8B-C). However, the specific molecular events responsible for this recruitment remain unclear. KIF2C might directly bind to PLK1, as previously suggested (Zhang *et al*., 2011). It might also recruit a partner, such as BRCA2, that binds specifically to PLK1. We observed that optogenetic KIF2C condensates co-localize with activated PLK1-pT210, whereas, at the spindle poles, they colocalize with PLK1 and BRCA2-pT207 (even after Aurora B inhibition), but not PLK1-pT210 (Suppl. Fig. 8B). Thus, colocalization of KIF2C and PLK1-pT210 is specific to condensates. When PLK1 is concentrated in the KIF2C condensates, it can phosphorylate KIF2C and its partners. In particular, it was reported that PLK1 phosphorylates S715 of KIF2C in mitosis (Shao *et al*., 2015). A lack of S715 phosphorylation impairs co-localization with BRCA2 phosphorylated at T207, as observed using the KIF2C S715A mutant (Figures 7D-E). We propose that condensates of KIF2C phosphorylated at S715 amplify the phosphorylation of BRCA2 by PLK1 at T207.

### KIF2C condensates locally amplify the activity of the PLK1 kinase

During mitosis, Aurora B is responsible for KIF2C localization at the centromere via the phosphorylation of Sgo2 (Andrews *et al*., 2004; Tanno *et al*., 2010). It also phosphorylates the large disordered region of KIF2C, which inhibits its depolymerization activity (Andrews *et al*., 2004), but activates PLK1 through phosphorylation of T210 in the activation loop of the kinase. PLK1 in turn phosphorylates KIF2C on S715, thus promoting its microtubule depolymerase activity. The multiple feedback loops involving Aurora B and PLK1 are essential to ensure the timely correction of aberrant kinetochore-microtubule attachments and a proper chromosome segregation (Joukov and De Nicolo, 2018). Here we used our artificial optogenetic system in order to decipher KIF2C condensate assembly mechanisms. We observed that, in absence of blue light, Aurora B is essential for KIF2C condensation (Figure 8E-F). As Aurora B activates PLK1, we propose that this activation is necessary for KIF2C condensate formation. However, exposure to blue light bypasses the requirement for active Aurora B, promoting KIF2C condensation even in the presence of Aurora B inhibitor (Figures 8E-F). As PLK1 is essential for light-induced condensate formation, we propose that optogenetic KIF2C condensates locally amplify PLK1 activity, thus bypassing Aurora B activity.

### KIF2C depolymerase activity is restrained to the periphery of the condensates

We observed that the catalytic activity of KIF2C facilitates its condensation in Noc (−) conditions (Figure 6B-C; Suppl. Fig. 6B-C). Moreover, KIF2C condensation in turn might regulate its depolymerase activity. Indeed, KIF2C condensates contain PLK1 that phosphorylates KIF2C S715, and this phosphorylation increases KIF2C affinity for microtubules, thus enhancing its microtubule depolymerase activity (Shao *et al*., 2015; Zhang *et al*., 2011). Also, tubulin is excluded from KIF2C condensates (Figure 7F). This is particularly clear after addition of nocodazole, because tubulin staining is mainly diffuse, and the few tubulin foci are all located at the surface of KIF2C foci. Consistently, it was previously reported that, in prometaphase chromosomes, KIF2C and tubulin localizations are close but distinct (Andrews *et al*., 2004). We observed that, in mitotic cells in Noc (−) conditions, KIF2C foci are located at the surface of the microtubules, in particular at their extremities (Figure 7G; Movie S2). It was also shown that KIF2C is enriched at the plus ends of polymerizing microtubules (Moore et al., 2005). Finally, SIM images revealed that KIF2C condensates are located next to CREST-marked centromeres (Figure 7H). The localization of KIF2C was proposed to be dynamic next to the centromeres, as tension develops across sister kinetochores (Andrews *et al*., 2004; Carmena *et al*., 2012). Our data suggests that KIF2C condensates form in proximity to both microtubules and kinetochores, and that the enzymatic activity of KIF2C is regulated through condensation; indeed, it may be enhanced by condensation, and as it can take place only at the periphery of KIF2C condensates.

## CONCLUSION

Understanding the temporal regulation of the key mitotic regulator PLK1 is essential, as PLK1 is an important oncogene in cancer initiation, progression and drug resistance. Here, we found that the microtubule depolymerase KIF2C, which facilitates proper microtubule-kinetochore attachment, forms nuclear condensates in mitosis. We further took advantage of a recently published optogenetic system to enhance the assembly of KIF2C biomolecular condensates in a time-controlled manner. We demonstrated that KIF2C condensates are membrane-less organelles that both fuse and grow from surrounding KIF2C molecules. Condensation depends on the N-terminal phospho-peptide binding domain of KIF2C. It is facilitated by PLK1-dependent interactions involving KIF2C and its partners. KIF2C condensates concentrate PLK1 and co-localize with BRCA2 phosphorylated at T207, which suggests that they favor BRCA2 phosphorylation by PLK1. This might in turn enhance the partition of PLK1 within KIF2C condensates, as a BRCA2 peptide centered at T207 directly binds to the Polo-Box Domain of PLK1. Moreover, KIF2C depolymerase activity promotes KIF2C condensate formation, and phosphorylation of KIF2C by PLK1 enhances its depolymerase activity. We propose that the assembly of KIF2C condensates locally amplifies both PLK1 and KIF2C catalytic activities, thus controlling kinetochore-microtubule attachment and favoring proper chromosome segregation during mitosis.

## Acknowledgements/fundings

We thank Emile Alghoul for the first KIF2C clonings. We thank Amandine Bonnet from CIGEX for her help in the production of the additional constructs for expression in mammalian cells. We thank Raphaël Ceccaldi for providing reagents. We thank V. Ropars for protein expression tests in insect cells. This work was supported by the CNRS and the CEA-Saclay, by the French Infrastructure for Integrated Structural Biology (https://frisbi.eu/, grant number ANR-10-INSB-05-01, Acronym FRISBI), by INSERM (PCSI 2022 grant BRCAPS coordinated by SZJ), by ANR (grant number ANR-21-CEA13-0030 coordinated by AC) and by ARC (ARC 2021 PJA3 grant N°248989 coordinated by SZJ). A.S. was funded by CEA, Synchrotron SOLEIL and FRM (grant N° FDT202304016927). R.C. was supported by a PSL University Fellowship. The Constantinou lab was supported by the French National Cancer Institute INCa (PLBIO 2021), by the French Agence Nationale de la Recherche ANR (AAPG2021 and AAPG2023), and by the Fondation MSD AVENIR. This work has benefited from the Imagerie-Gif core facility supported by l’Agence Nationale de la Recherche (ANR-11-EQPX-0029/Morphoscope, ANR-10-INBS-04/FranceBioImaging; ANR-11-IDEX-0003-02/ Saclay Plant Sciences). Financial support from the IR INFRANALYTICS FR2054 for conducting the research is also gratefully acknowledged.

## Author contributions

A.S., M.J., S.M., T.E., R.C., G.B., R.G., D.B. conceived and performed the experiments. S.J., S.M., F.X.T., R.L.B., Au.C., J.B., S.Z.J. conceived and supervised the experiments. A.S., M.J., G.B., J.B., S.Z.J analyzed the experiments. S.J., R.L.B., An.C. provided expertise and feedback. C.F. provided reagents. A.S., S.Z.J. wrote the original draft. S.J., R.L.B., F.X.T. reviewed and edited the article. Au.C., An.C, S.Z.J. provided funding.

## Supplemental information titles and legends

**Suppl. Figure 1. Extended data for Fig. 1.**
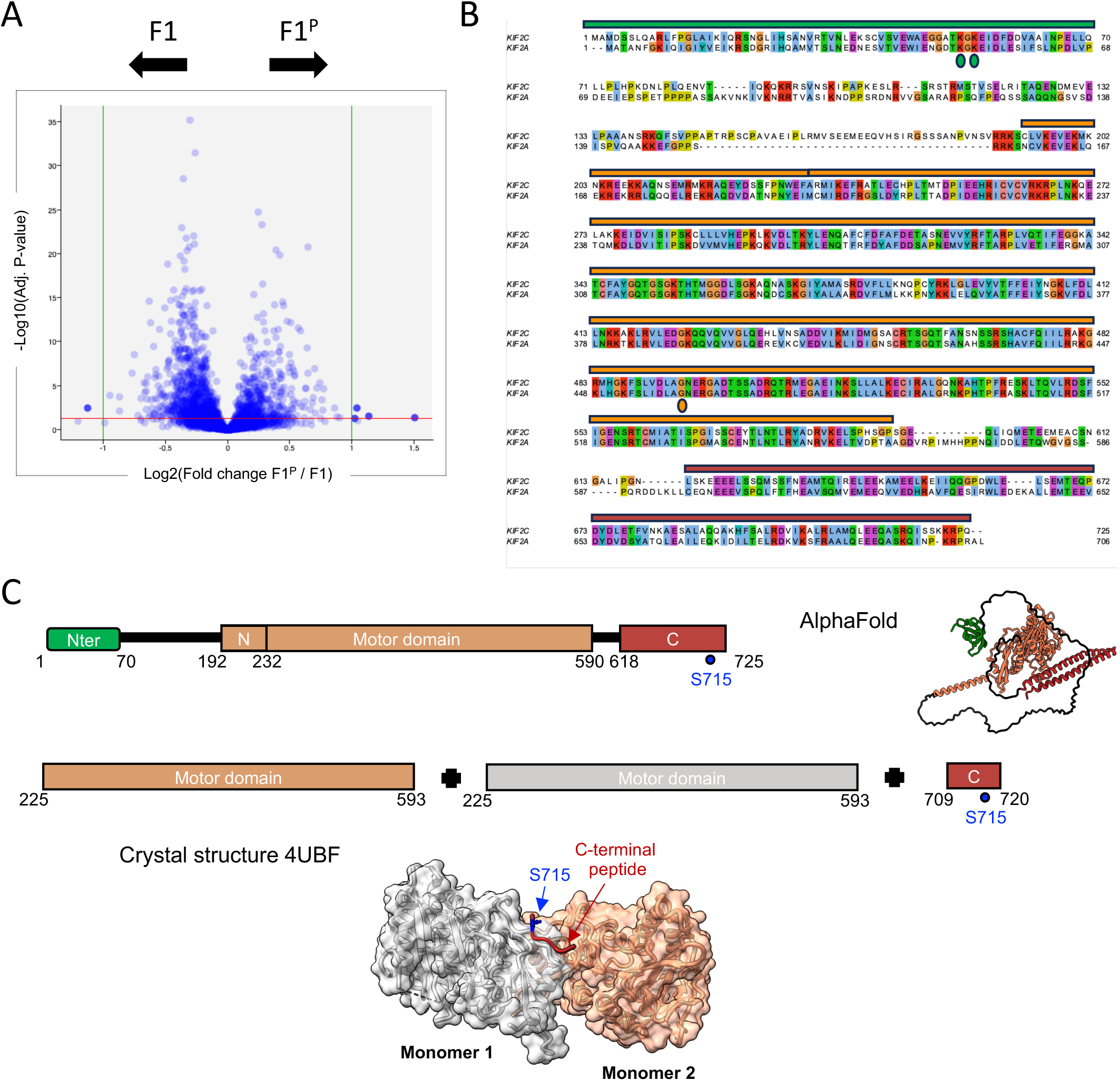
(A) Extended data for Fig. 1B. Volcano plots showing proteins from asynchronous HeLa cell extracts that are identified by mass spectrometry as binding to (left) BRCA2_167-270_ phosphorylated by PLK1 (here named F1^P^) versus non phosphorylated BRCA2_167-270_ (here named F1). (B) Extended data for Fig. 6C. Alignment of the sequences of human KIFC and KI2A, with residues of KIF2C mutated in this study indicated by colored circles. Bars above the sequences correspond to the domain as shown in Figure 1C. (C) Dimerisation of KIF2C, as observed by X-ray crystalllography. The domain organisation of KIF2C, and its AlphaFold model, are presented in the upper panel, while the crystallized domains, and the associated dimeric KIF2C structure, are presented in the lower panel (PDB code: 4UBF; Talapatra et al., Elife 2015). The two motor domain monomers are colored in orange and grey, respectively. The C-terminal peptide is displayed in red sticks, with S715 in blue. Mutation S715E disrupts binding of the peptide to the motor domains (Talapatra et al., Elife 2015).

**Suppl. Figure 2. Extended data for Fig. 2 and 3B.**
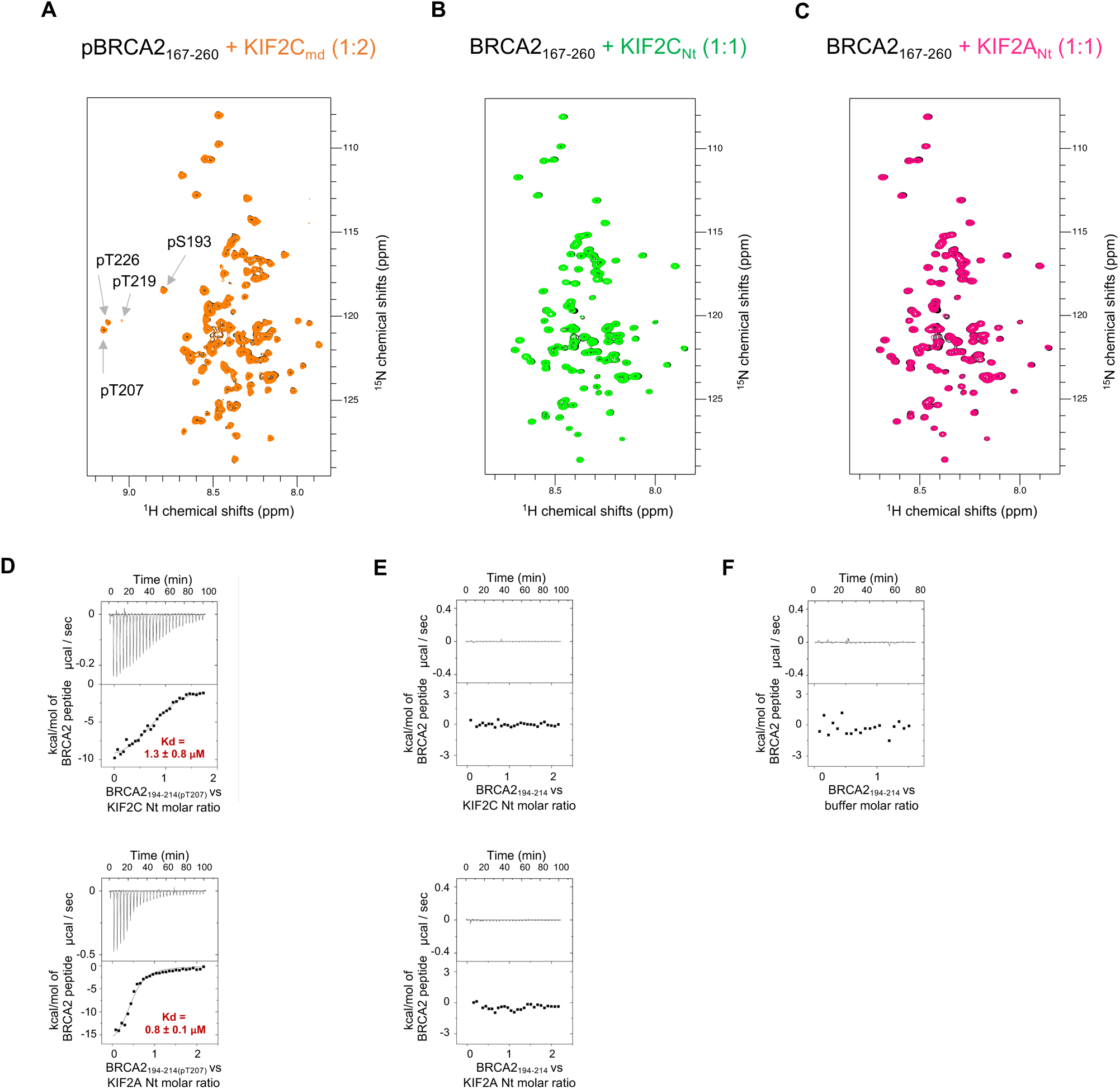
(A) Extended data for Fig. 2A. Superposition of the 2D NMR ^1^H-^15^N SO-FAST HMQC spectra of phosphorylated BRCA2_167-270_ (50 μM) in the absence (black) and presence (orange) of the neck and motor domain of KIF2C (KIF2C_md_, 100 μM) recorded at 600 MHz and 283 K. (B) Superposition of the 2D NMR ^1^H-^15^N SO-FAST HMQC spectra of non phosphorylated BRCA2_167-270_ in the absence (black) and presence (green) of the N-terminal domain of KIF2C recorded at 700 MHz and 283 K. (C) Superposition of the 2D NMR ^1^H-^15^N SO-FAST HMQC spectra of non phosphorylated BRCA2_167-270_ in the absence (black) and presence (pink) of the N-terminal domain of KIF2A recorded at 700 MHz and 283 K. (D) Duplicates for Fig. 3B, showing binding of either KIF2C Nt or KIF2A Nt to BRCA2_194-214(pT207)_. (E) ITC analyses indicating that both KIF2C Nt and KIF2A Nt do not bind to non-phosphorylated BRCA2_194-214_. (F) ITC analysis indicating that the dilution of the peptide BRCA2_194-214(pT207)_ in the buffer does not generate a heat signal.

**Suppl. Figure 3. Extended data for Fig. 3C-E.**
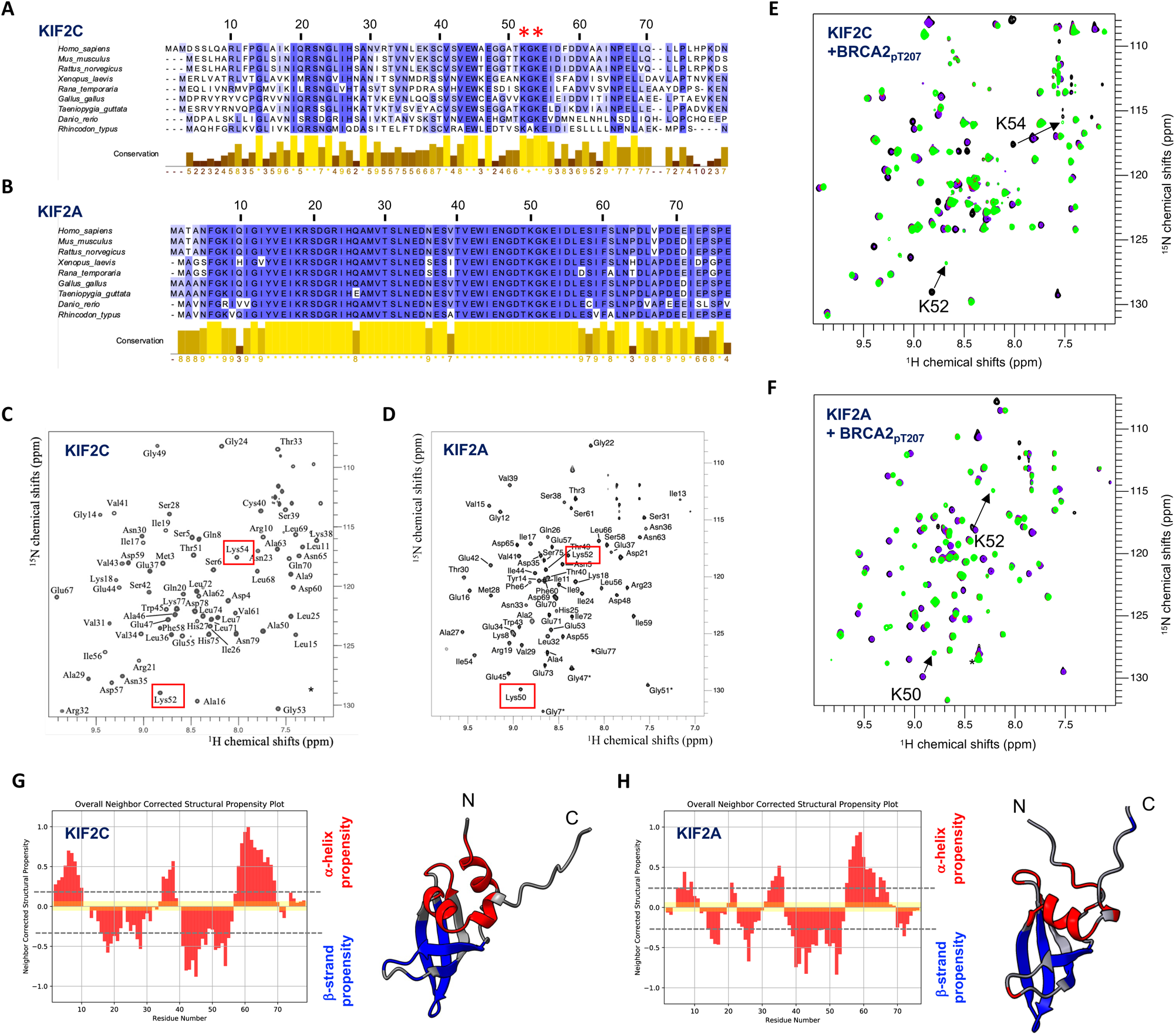
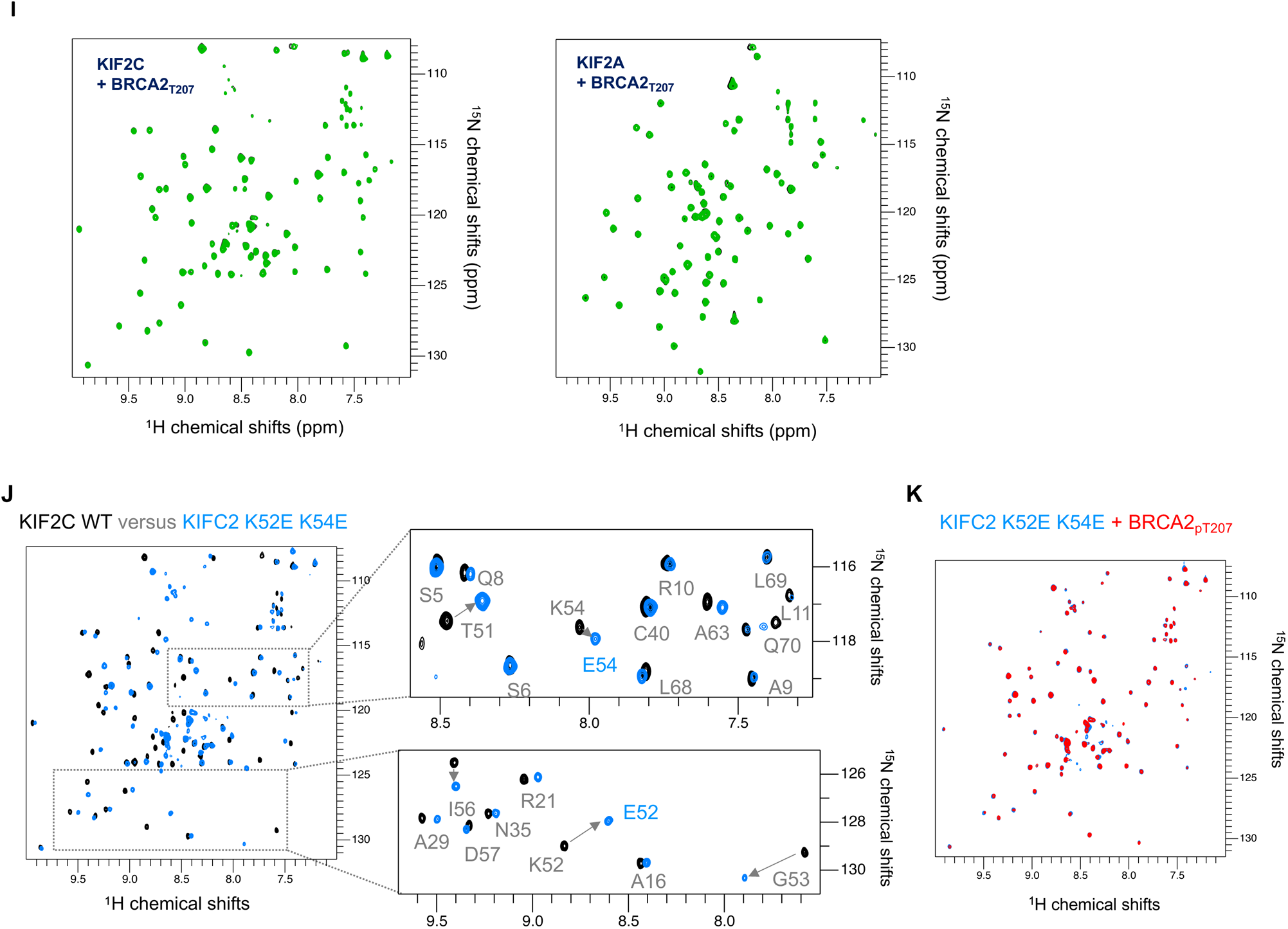
(A,B) Extended data for Fig. 3E. Sequence alignement of the N-terminal domains of (A) KIF2C and (B) KIF2A, from human to fishes, as displayed using Jalview. (C,D) Extended data for Fig. 3C-D. 2D NMR ^1^H-^15^N HSQC spectrum of the N-terminal domain of KIF2C and KIF2A. Residues corresponding to each peak are marked. Stars refer to peaks that are shifted from one ^15^N spectral width. (E,F) Extended data for Fig. 3C-D. Superimposition of the 2D NMR ^1^H-^15^N SO-FAST HMQC spectra of ^15^N labeled (E) KIF2C and (F) KIF2A N-terminal domains either free (black) or in the presence of 0.5 (mauve), 1.0 (red) or 2.0 (green) equivalent of BRCA2_194-214(pT207)_. (G,H) Extended data for Fig. 3C-D. Neighbor Corrected Structural Propensity as a function of the residue number, as deduced from (G) KIF2C and (H) KIF2A ^1^Hn, ^15^N, Cα, Cβ and Co chemical shifts (website: https://st-protein02.chem.au.dk/ncSPC/cgi-bin/selection_screen_ncSPC.py), and (G) KIF2C and (H) KIF2A N-terminal domain models (AlphaFold entries Q99661 and O00139) colored as a function of the residue chemical shift analysis: residues that show a tendency (propensity > 0.25) to form α-helix and β-strand are colored in red and blue, respectively. (I) Extended data for Fig. 3C-D. Superimposition of the 2D NMR ^1^H-^15^N SO-FAST HMQC spectra of ^15^N labeled KIF2C and KIF2A N-terminal domains either free (black) or in the presence of 2.0 (green) equivalent of BRCA2_194-214_. (J) Superposition of the 2D NMR ^1^H-^15^N HSQC spectra of the N-terminal domain of KIF2C either WT (black) or mutated (blue), and zooms showing the peaks corresponding to the mutations K52E and K54E. (K) Superposition of the 2D NMR ^1^H-^15^N SO-FAST HMQC spectra of the N-terminal domain of KIF2C mutant K52E K54E, either free (blue) or in the presence of a BRCA2_194-214(pT207)_ (red).

**Suppl. Figure 4. Extended data for Fig. 4.**
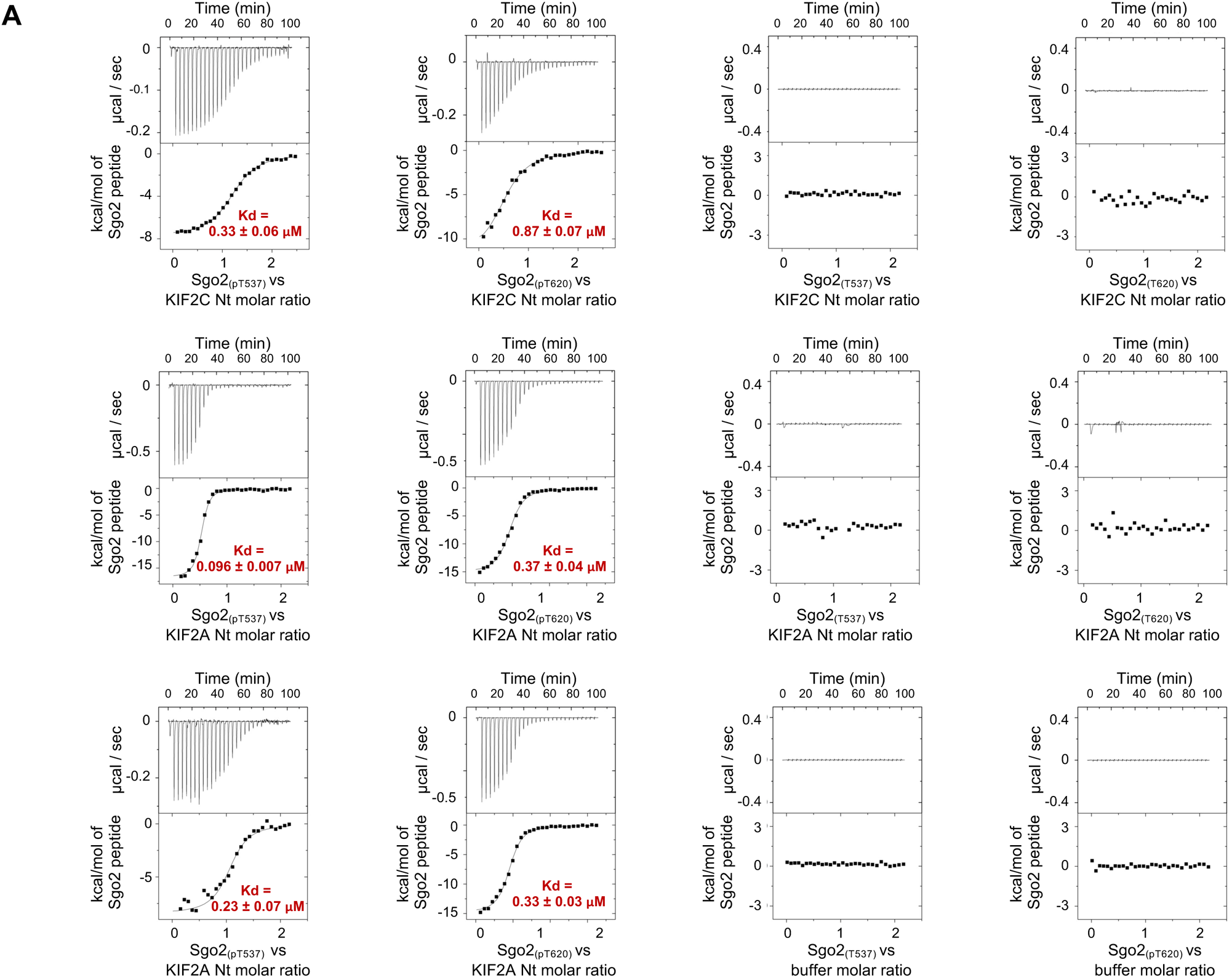
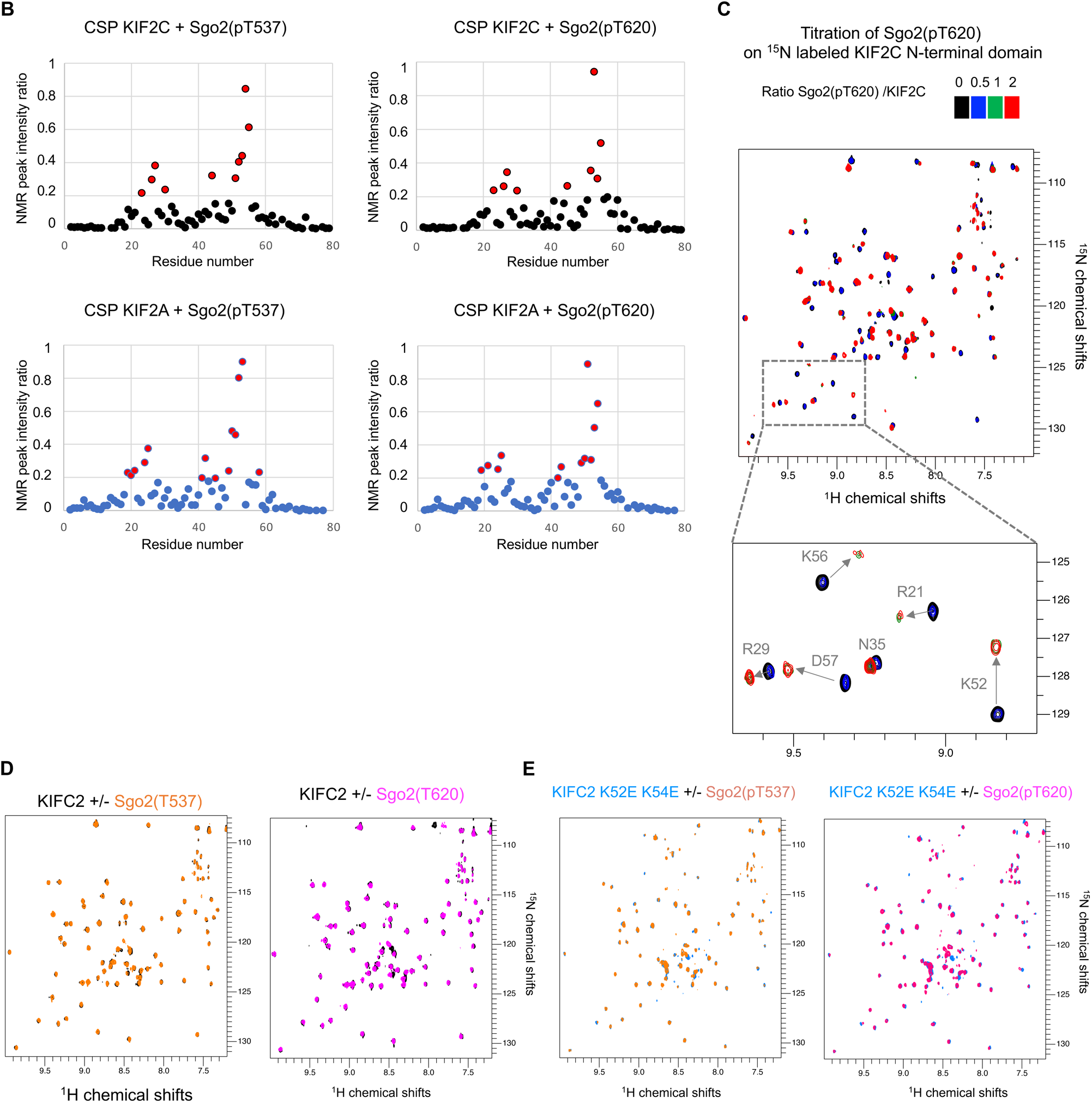
(A) Extended data for Fig. 4C. Duplicates and control ITC experiments. (B) Extended data for Fig. 4D. NMR chemical shift perturbation (CSP) analyses of the titrations of ^15^N labeled KIF2C or KIF2A N-terminal domain with Sgo2 peptide phosphorylated at T537 or T620. Values larger than 0.2 ppm are marked in red, and the spatial distribution of the corresponding residues is displayed in Figure 4D. (C) Extended data for Fig. 4D. Superposition of the 2D NMR ^1^H-^15^N SO-FAST HMQC spectra of the N-terminal domain of KIF2C in the presence of increasing concentrations of Sgo2_(pT620)_, illustrating that peaks corresponding to free and bound states are decreasing and increasing upon addition of the peptide, respectively. The weak intensity of several peaks corresponding to the bound state indicates that the exchange between the free and bound states is intermediate to slow on the NMR time scale. (D) Extended data for Fig. 4D. Superposition of the 2D NMR ^1^H-^15^N SO-FAST HMQC spectra of the N-terminal domain of KIF2C in the absence (black) or presence of non-phosphorylated Sgo2_(T537)_ (orange) or Sgo2_(T620)_ (pink), illustrating that the kinesin domain does not bind to non-phosphorylated Sgo2 peptides. (E) Superposition of the 2D NMR ^1^H-^15^N SO-FAST HMQC spectra of the N-terminal domain of KIF2C K52E-K54E in the absence (blue) or presence of phosphorylated Sgo2_(pT537)_ (orange) or Sgo2_(pT620)_ (pink), showing that the mutated domain does not bind to non-phosphorylated Sgo2 peptides.

**Suppl. Figure 5. Extended data for Fig. 5.**
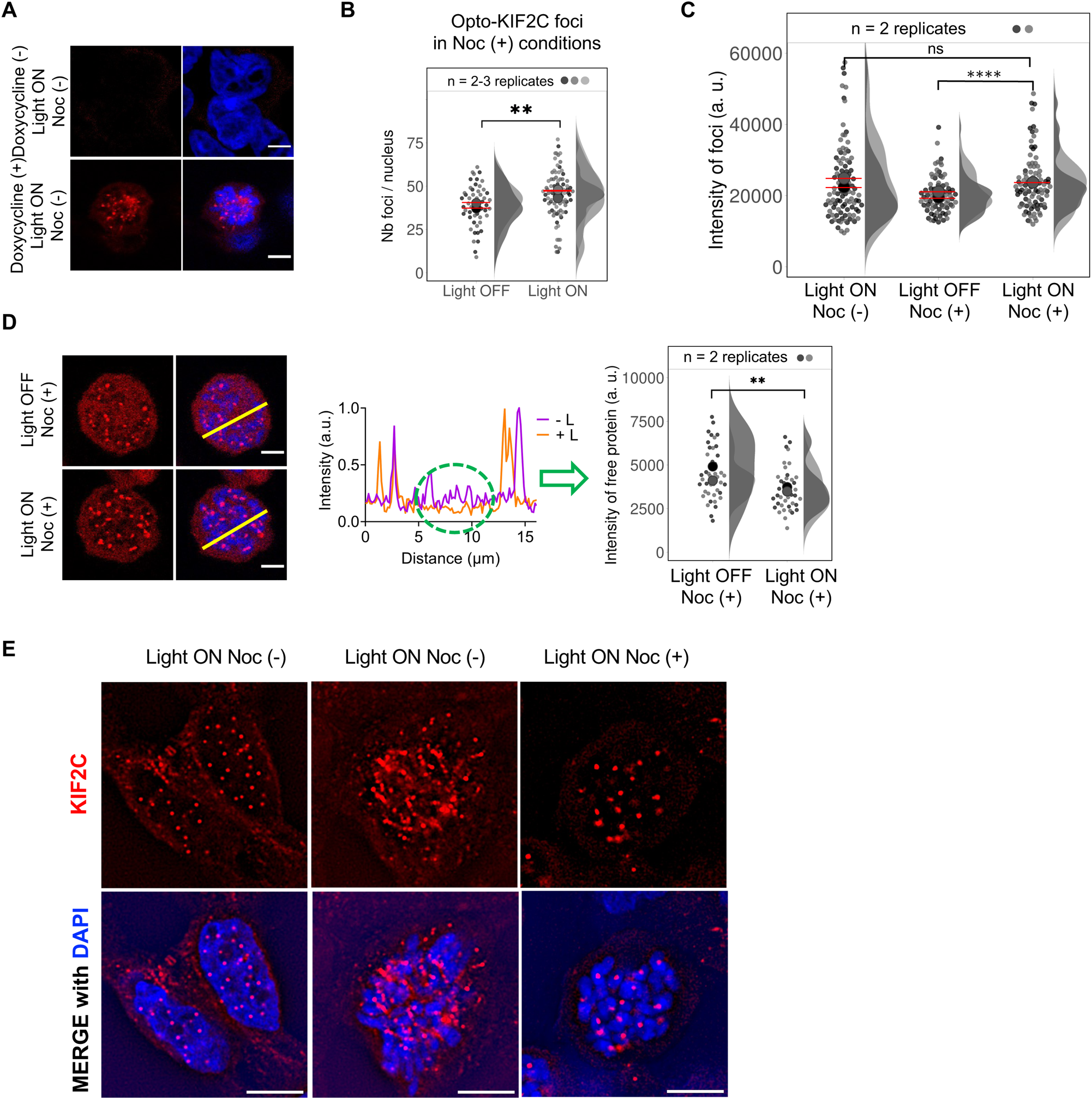
(A) Extended data for Fig. 5B. Representative fluorescence images obtained before (upper line) and after (lower line) induction by doxycycline of the expression of Opto-KIF2C WT and illumination of the cells with blue light (light ON). Opto-KIF2C foci are observed only upon addition of doxycycline. DNA is stained with Hoechst 33258. Scale bars: 5 µm. (B) Quantification of the number of Opto-KIF2C foci per cell as measured on the images recorded as in Fig. 5B (lines 2, 4). P-value is 0.0026. (C) Measurement of the fluorescence intensity of foci in different light and nocodazole conditions. The p-value calculated between the distributions of foci intensities measured under light OFF / Noc (+) and light ON / Noc (+) conditions is lower than 0.0001, whereas the p-value calculated between the distributions of intensities measured under light ON / Noc (−) and light ON / Noc (+) conditions is 0.4639. (D) Representative fluorescence images obtained after induction by doxycycline and addition of nocodazole of the expression of Opto-KIF2C WT before (upper line) and after (lower line) illumination of the cells with blue light. Scale bars: 5 µm. Line scans show a decrease in free protein in the nucleus after 5 min of illumination. This decrease is quantified in the plot on the right. The p-value associated to this plot is 0.0014. (E) Extended data for Fig. 5C. SIM images obtained with the Opto-KIF2C construct in light ON conditions and either without (Noc (−)) or with (Noc (+)) nocodazole. Scale bars: 5 µm.

**Suppl. Figure 6. Extended data for Fig. 6.**
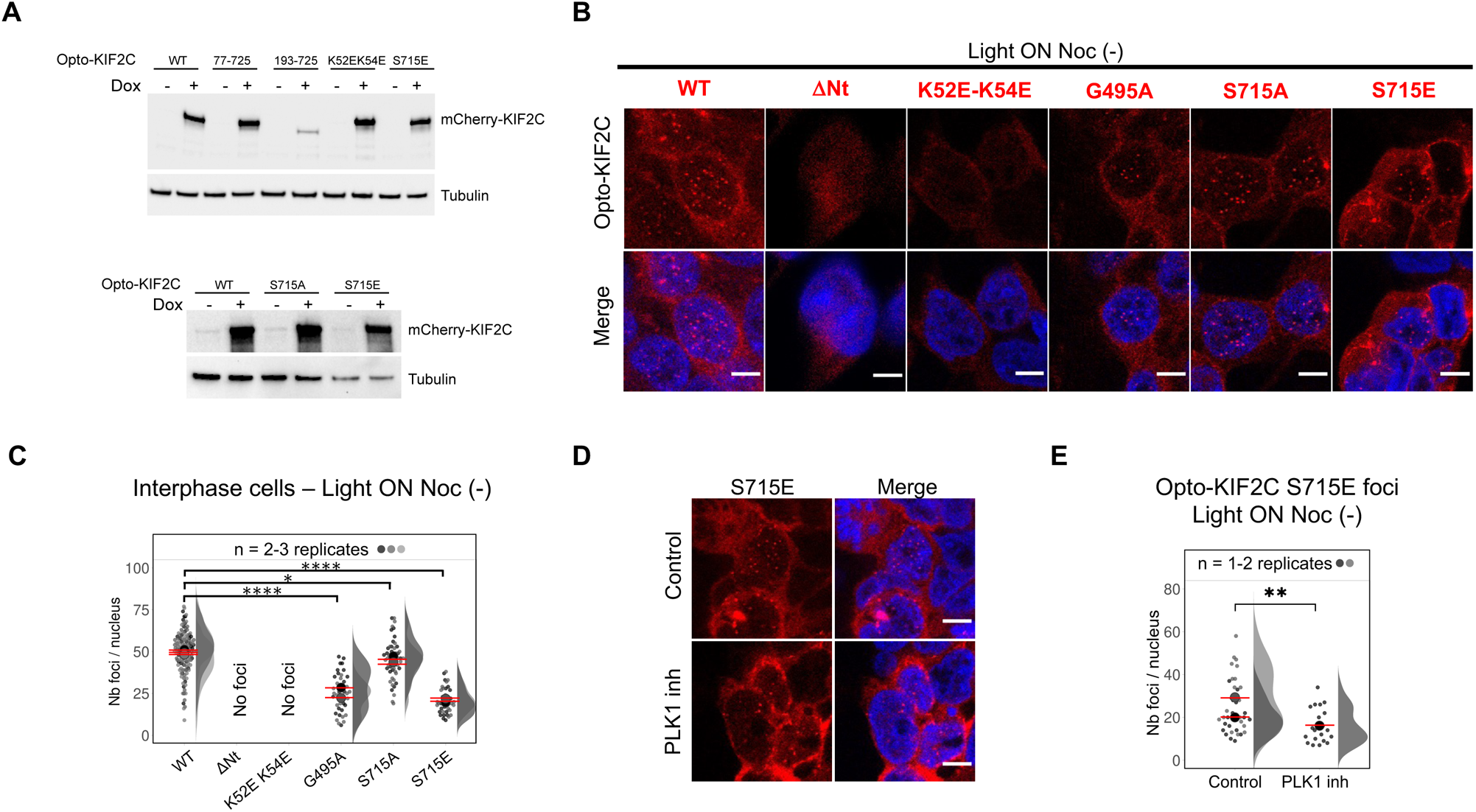
(A) Extended data for Fig. 6A, showing the expression of Opto-KIF2C mutants in cells. In mutant 77-725, the N-terminal domain of KIF2C is deleted. In mutant 193-725, the N-terminal domain and the disordered region of KIF2C are deleted; this mutant is poorly expressed in cells and was not further used in this study. In mutant K52EK54E, the two critical lysines of the N-terminal domain of KIF2C are mutated into glutamic acid. In the last two mutants, S715 is mutated into either alanine or the phosphomimetic glutamic acid. (B) Extended data for Fig. 6B. Representative fluorescence images obtained after induction by doxycycline of the expression of Opto-KIF2C mutants and illumination of the interphase cells with blue light (light ON). Scale bars: 5 µm. (C) Quantification of the amount of foci per cell as measured on the images recorded as in panel (B). P-values corresponding to G495A and S715E are lower than 0.0001, whereas that of S715A is 0.0105. (D) Extended data for Fig. 6D. Impact of PLK1 inhibition on KIF2C S715E foci formation (Light ON Noc (−)). (E) Quantification of the amount of foci per cell as measured on the images recorded as in panel (D). The p-value measured between the distributions of numbers of foci observed in the absence and presence of PLK1 inhibitor is 0.0050.

**Suppl. Figure 7. Extended data for Fig. 7.**
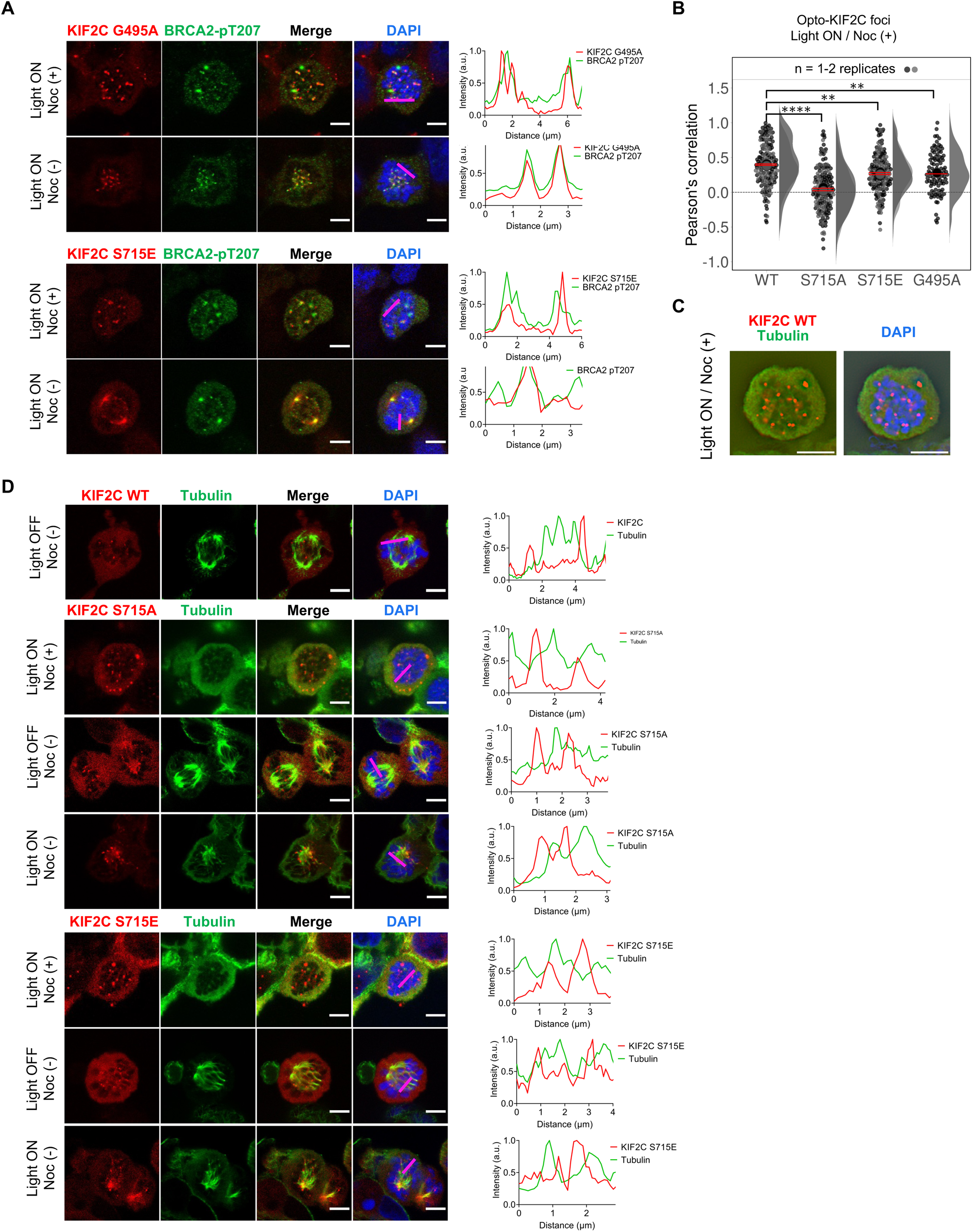
(A) Extended data for Fig. 7D. Representative immunofluorescence images of Opto-KIF2C G495A and S715E (red channel), BRCA2-pT207 (green channel) and DNA (blue channel). Line scans on the right show co-localization. (B) Extended data for Fig. 7E. Quantification, using Pearson’s correlation coefficient, of the co-localization between Opto-KIF2C (WT and mutants) and BRCA2-pT207 after addition of nocodazole (Noc (+)). Median shown in red. P-values corresponding to G495A and S715E are 0.0025 and 0.0034, respectively, and the p-value corresponding to S715A is lower than 0.0001. (C) Extended data for Fig. 7G. SIM images obtained with Opto-KIF2C (red), tubulin (green) and DAPI (blue) in light ON and Noc (+) conditions. (D) Extended data for Fig. 7F. Representative immunofluorescence images of Opto-KIF2C WT, S715A and S715E (red channel), tubulin (green channel) and DNA (blue channel). Line scans on the right show co-localization. Scale bars: 5 µm.

**Suppl. Figure 8. Extended data for Fig. 8.**
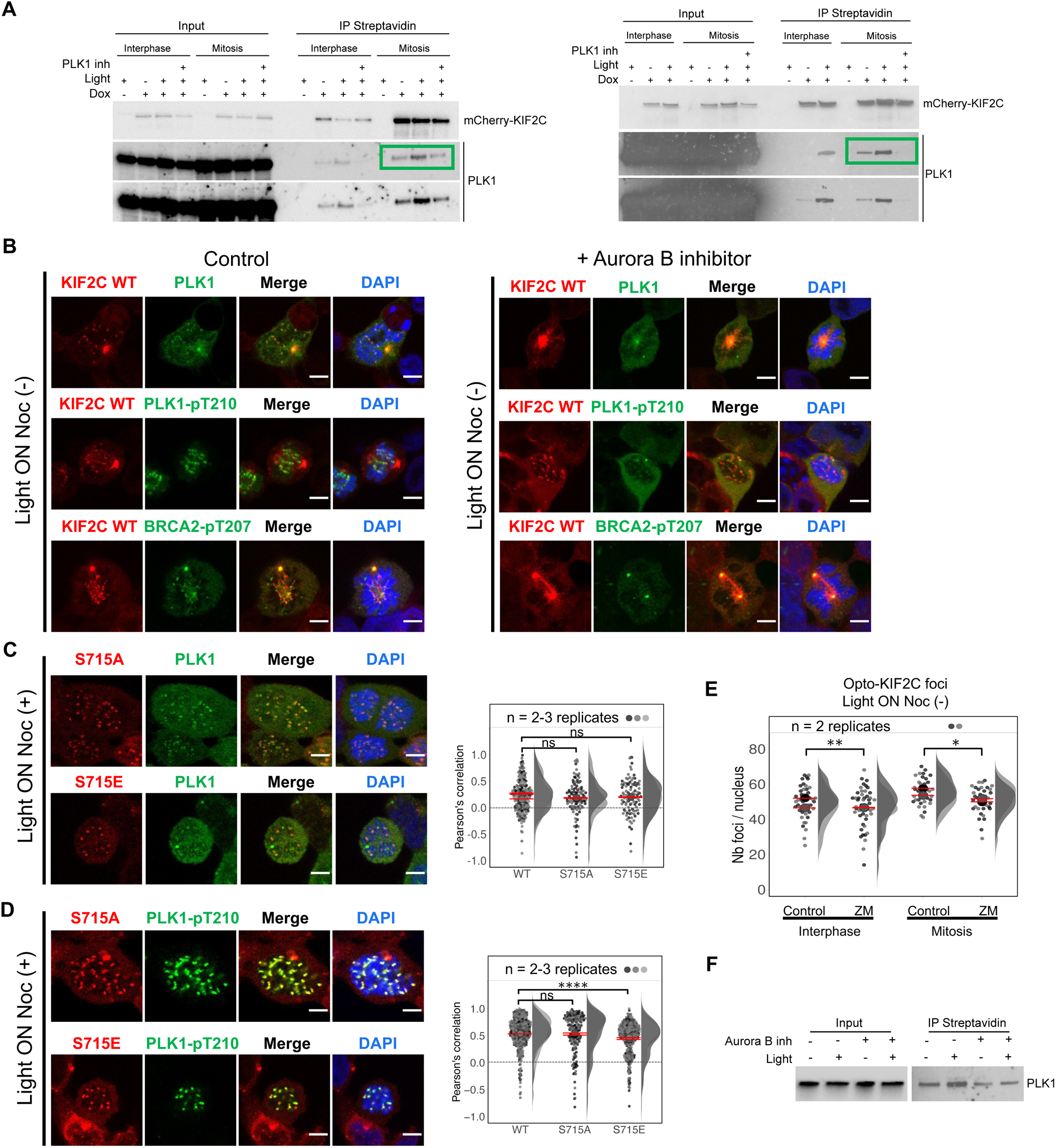
(A) Two replicates of Figure 8A: PLK1 was identified as enriched in KIF2C condensates. (B) Extended data for Fig. 8C-D. Representative immunofluorescence images of Opto-KIF2C WT (red channel), PLK1, PLK1-pT210 or BRCA2-pT207 (green channel) and DNA (blue channel) in light ON and Noc (−) conditions. Images were obtained without (left) and upon addition of (right) the Aurora B inhibitor ZM447439, respectively. (C) Extended data for Fig. 8C-D. Representative immunofluorescence images of Opto-KIF2C S715A and S715E, PLK1 (green channel) and DNA (blue channel), and associated quantification of the colocalisation of Opto-KIF2C and PLK1, in light ON and Noc (+) conditions. P-values corresponding to S715A and S715E are 0.3983 and 0.6972, respectively. (D) Extended data for Fig. 8C-D. Representative immunofluorescence images of Opto-KIF2C S715A and S715E, PLK1-pT210 (green channel) and DNA (blue channel), and associated quantification of the colocalisation of Opto-KIF2C and PLK1-pT210, in light ON and Noc (+) conditions. P-values corresponding to S715A and S715E are 0.3286 and lower than 0.001, respectively. (E) Quantification of the number of Opto-KIF2C foci per nucleus from the analysis of immunofluorescence images recorded in light ON and Noc (−) conditions, in the absence and presence of the Aurora B inhibitor ZM447439, as presented in panel (B). P-values are 0.0059 and 0.0169 in interphase and mitotic cells, respectively. (F) Identification of Turbo-Opto-KIF2C partners by Western Blot. PLK1 was enriched upon illumination and decreased upon addition of the Aurora B inhibitor. Scale bars: 5 µm.

**Supplementary Table 1:**
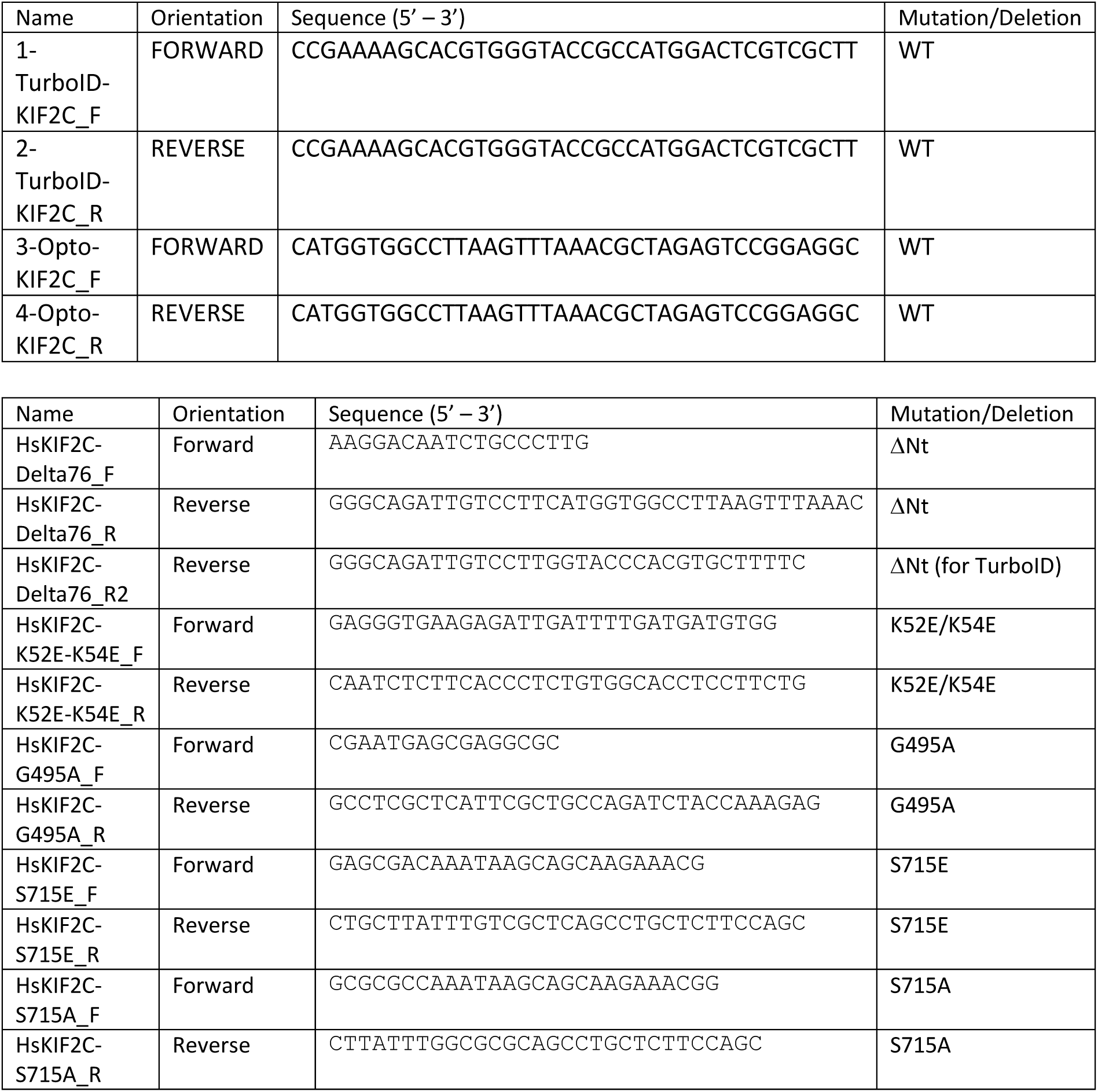
Oligonucleotides used for plasmid construction.

**Supplementary Table 2:** Opto-constructions of KIF2C WT and mutants.

| Construct | Mutation |
| --- | --- |
| pcDNA5-FRT-Hygro-HsKIF2C-mCherry-AtCRY2-1-498 | WT |
| pcDNA5-FRT-Hygro-HsKIF2C-77-725-mCherry-AtCRY2-1-498 | ΔNt |
| pcDNA5-FRT-Hygro-HsKIF2C-1-725 –K52E-K54E-mCherry-AtCRY2-1-498 | K52E/K54E |
| pcDNA5-FRT-Hygro-HsKIF2C-1-725–V145A-G495A-mCherry-AtCRY2-1-498 | G495A |
| pcDNA5-FRT-Hygro-HsKIF2C-1-725–V145A-S715E-mCherry-AtCRY2-1-498 | S715E |
| pcDNA5-FRT-Hygro-HsKIF2C-1-725–V145A-S715A-mCherry-AtCRY2-1-498 | S715A |
| pcDNA5-FRT-Hygro-TurboID-HsKIF2C-mCherry-AtCRY2-1-498 | WT |
| pcDNA5-FRT-Hygro-TurboID-HsKIF2C-77-725-mCherry-AtCRY2-1-498 | ΔNt |
| pcDNA5-FRT-Hygro-TurboID-HsKIF2C-2-725-K52E-K54E-mCherry-AtCRY2-1-498 | K52E K54E |
| pcDNA5-FRT-Hygro-TurboID-HsKIF2C-2-725-G495A-mCherry-AtCRY2-1-498 | G495A |
| pcDNA5-FRT-Hygro-TurboID-HsKIF2C-2-725-S715E-mCherry-AtCRY2-1-498 | S715E |
| pcDNA5-FRT-Hygro-TurboID-HsKIF2C-2-725-S715A-mCherry-AtCRY2-1-498 | S715A |

**Supplementary Table 3:** Synthetic peptide sequences for NMR and ITC.

| Name | Sequence | Supplier |
| --- | --- | --- |
| BRCA2<sub>194-214</sub> | WSSSLATPPTLSSTVLIVRNE | GenScript |
| BRCA2<sub>194-214</sub>(pT207) | WSSSLATPPTLSS(pT)VLIVRNE | GenScript |
| Sgo2<sub>531-544</sub> | SKASRQTFVIHKLE | GenScript |
| Sgo2<sub>531-544</sub>(pT537) | SKASRQ(pT)FVIHKLE | GenScript |
| Sgo2<sub>614-627</sub> | KKVNRKTEIISGMN | GenScript |
| Sgo2<sub>614-627</sub>(pT620) | KKVNRK(pT)EIISGMN | GenScript |

**Supplementary Table 4:** List of antibodies used for IF and WB.

| Antibodies | Source | Identifier |
| --- | --- | --- |
| Anti-PLK1 (phospho T210) antibody [2A3] | Abcam | Cat# ab39068 |
| Anti-PLK1 | Sigma | Cat# 05-844 |
| Donkey anti-rabbit Alexa-488 | Molecular Probes | Cat# A-21206 |
| Donkey anti-mouse Alexa-488 | Thermo Fisher Scientific | Cat# A-21202 |
| Donkey anti-rabbit Alexa-594 | Thermo Fisher Scientific | Cat# A-21207 |
| Goat anti-human Alexa-555 | Thermo Fisher Scientific | Cat# 10729014/A21433 |
| Mouse anti-KIF2C | Santa Cruz Biotechnology | Cat# sc-81305 |
| Human anti-Crest | Antibodies Incorporated | Cat#15-234-0001 |
| Anti-mouse-BRCA2 | EMD Millipore | Cat# D54931 |
| Anti-α-Tubulin, clone DM1A | EMD Millipore | Cat# MABT205 |
| Anti-rabbit-pBRCA2-T207 | Genscript |  |
| Anti-mouse IgG, HRP linked Antibody | Cell Signaling Technology | Cat# 7076; RRID:AB_330924 |
| Anti-rabbit IgG, HRP linked Antibody | Cell Signaling Technology | Cat# 7074; RRID:AB_2099233 |

**Supplementary Movie 1:** Time-lapse microscopy movie of activated opto-KIF2C condensates that are fusing (light ON, Noc (−)).

**Supplementary Movie 2:** 3D SIM image obtained with Opto-KIF2C (red), tubulin (green) and DAPI (blue) in light ON and Noc (−) conditions.

## STAR Methods

### Cell lines and synchronization

Flp-In T-REx 293 3 cell lines were grown under standard conditions (37°C, 5% CO2) in Dulbecco’s modified Eagle’s medium (Merck-Sigma-Aldrich, D5796) containing 10% Fetal Bovine Serum (FBS) and penicillin-streptomycin. Parental cells were selected with 100mg/ml Zeocin and Flp-In 293 T-REx derived stable cell lines were maintained with 5mg/ml Blasticidin and 50mg/ml Hygromycin B. DLD1+/+ cells were grown in RPMI media supplemented with 2 mM L-Glutamine (prepared in the culture facility of CBMSO) and 10% Fetal Bovine Serum (FBS) (BioWest S1810-500). DLD1 cells stably expressing EGFP-MBP-BRCA2 WT were grown in RPMI media supplemented with 2 mM L-Glutamine, 10% FBS, 1 mg/ml G418 (Enzo Life Sciences ALX-380-013-G005) and 0.1 mg/ml Hygromycin B (Thermo Scientific 10687010). All cells were grown in a humidified incubator at 37°C and 5% CO_2_. For synchronization of cells in mitosis, nocodazole (100–300 ng/ml, Sigma-Aldrich) was added to the growth media and the cells were cultured for 14 h before harvesting.

### Plasmids

Oligonucleotides used for plasmid construction are listed in Suppl. Table S1. For expression in bacteria, optimized genes coding for human 6His-AviTag-BRCA2_167-260_ and 8His-TEVsite-BRCA2_48–218_ were synthetized by Genscript and cloned in a pETM13 vector. In addition, optimized genes coding for human 8His-GB1-TEVsite-KIF2C_1-79_, its K52E/K54E mutant, and 8His-GB1-TEVsite-KIF2A_1-77_ were synthetized by Genscript and cloned in a pET-22b vector. For expression in mammalian cells, we first obtained pcDNA5-FRT-Hygro plasmids coding for HsKIF2C-2-725-V145A-mCherry-AtCRY2-1-498 and TurboID-HsKIF2C-2-725-V145A-mCherry-AtCRY2-1-498. These plasmids were mutated to generate the list of constructs displayed in Suppl. Table S2. The EGFP-MBP-tagged BRCA2 cDNA was previously described (Ehlen *et al*., 2020).

### Cloning

Primers used for plasmid construction are listed in the KRT. For pcDNA5_FRT_TurboID constructs, KIF2C cDNA was amplified with primers 1 and 2 (See supplementary Table 1) using Phusion High-Fidelity DNA Polymerase. The amplified sequence was inserted into the KpnI site of pCDNA5_FRT_TO_TurboID-mCherry-Cry2 (Addgene 166504). pcDNA5_FRT_opto-KIF2C-cDNA was obtained by removing the TurboID from the pcDNA5_FRT_TurboID-opto-KIF2C construct using the primer 3 and 4. Mutations in KIF2C were generated using the QuickChangeMulti Site-Directed Mutagenesis Kit with the primers listed in Supplementary Table 1.

### Generation of stable cell lines

Flp-In 293 T-REx cells are seeded to reach 80-90% confluence on the day of transfection. pcDNA5_FRT_TO expression plasmids were mixed with pOG44 encoding the Flp recombinase at a 1:7 ratio in opti-MEM. For a single transfection in a 6 well plate, 500ng of the expression plasmid was mixed with 3.5mg of pOG44 in 250uL opti-MEM. Additionally, 8uL Lipofectamine 2000 Transfection Reagent was added to 250uL opti-MEM. After an incubation period of 5 minutes at room temperature, both solutions were mixed and incubated for a further 15 minutes at room temperature. The mixture was then pipetted dropwise onto the cells. The medium was changed after 6 hours. At 48 hours post-transfection, the cells were transferred to a 100mm petri dish, and 24 hours later, the selection was performed by adding 5mg/mL Blasticidin and 50mg/mL Hygromycin B. Clones were pooled, and the cells were examined for the expression of the construct by immunoblotting and fluorescence microscopy.

### Production of the recombinant Avi-tagged BRCA2_167-260_ peptide

A recombinant 6His-Avi-BRCA2_167-260_ construct was designed that contains the 15 amino acid sequence GLNDIFEAQKIEWHE, so-called Avi-Tag (Fairhead and Howarth, 2015). It was produced in *E. coli* BL21 (DE3) Star cells in either its unlabeled form for pulldown experiments or its ^15^N-labeled form for NMR experiments. Cells were grown in M9 minimal medium containing 0.5 g/l ^15^NH_4_Cl when ^15^N labeling was needed. The bacterial culture was induced at an OD_400_ of 0.6-0.8 with 1 mM IPTG and was further incubated during 4 h at 37° C. Harvested cells were resuspended in 50 mM Tris-HCl pH 8.0, 50 mM NaCl, 5 mM DTT, 2 mM EDTA, 1 mM PMSF, 1 mM ATP, 5 mM MgSO_4_, 500 µg of lysozyme, 0.5 µL of benzonase (E1014; Millipore). They were sonicated on ice 2.5 min in total with 1s ON/1s OFF cycle of sonication (50% amplitude), and the lysate was clarified by centrifugation during 15 min at 15,000 g at 4°C. The soluble fraction was loaded on a Ni-NTA poly-histidine-affinity column (5 mL HisTrap Excel, GE Healthcare) at a 2 mL/min flow rate. The column was washed with a solution containing 50 mM Tris-HCl at pH 8.0, 50 mM NaCl, 1 mM DTT. The sample was then eluted with an imidazole gradient over 45 mL, the buffer containing 50 mM Tris-HCl at pH 8.0, 50 mM NaCl, 1 M imidazole, 1 mM DTT. The sample was boiled 5 min at 95°C and spun down 5 min at 16,000 g at 4°C. It was concentrated using Novagen concentrators with 3.5 kDa cut off membranes centrifuged at 5,000 g and later injected on a gel filtration column (Superdex 16/600 HiLoad 75 pg) equilibrated with a Dulbecco’s phosphate buffered saline (DPBS) (D1408; Sigma) at pH 7.2. Fractions were pooled, 1 mM of fresh DTT was added and the sample was concentrated using a 3 kDa cut off concentrator centrifuged at 5,000 g. EDTA-free protease inhibitors (cOmplete EDTA-free, Sigma-Aldrich) were added at a final 1X concentration. The quality of the purified proteins was analyzed by SDS-PAGE and the protein concentrations were determined by spectrophotometry using the absorbance at 280 nm. All concentrated proteins were stored in small aliquots at −80 °C.

### Production of the recombinant BRCA2_48-218_ peptide

^15^N-labeled 8His-TEVsite-BRCA2_48-218_ was produced by transforming *E. coli* BL21 (DE3) Star cells, cultured in M9 minimal medium containing 0.5 g/l ^15^NH_4_Cl, and supplemented with 100 μg/ml kanamycin, at 37 °C. Expression was induced at an OD_400_ of 0.6-0.8 with 1 mM IPTG and cells were harvested after 3-4 h of incubation by centrifugation. The bacterial pellet was resuspended in 35 mL of a solution containing 50 mM Tris-HCl at pH 8.0, 50 mM NaCl, 5 mM DTT, 2 mM EDTA, 1 mM PMSF, 1 mM ATP, 5 mM MgSO_4_, 500 μg lysozyme, 0.5 μL benzonase (E1014; Millipore). Cells were sonicated on ice 2.5 min in total with 1s ON/1s OFF cycle of sonication (50% amplitude), and the lysate was clarified by centrifugation during 15 min at 15,000 g at 4°C. The soluble fraction was loaded on a Ni-NTA histidine-affinity column (5 mL HisTrap Excel, GE Healthcare) using a 2 mL/min flow rate. The column was washed with a solution containing 50 mM Tris HCl at pH 8.0, 50 mM NaCl, 1 mM DTT. The sample was eluted using an imidazole gradient over 45 mL, the final buffer containing 50 mM Tris HCl at pH 8.0, 50 mM NaCl, 1 M imidazole, 1 mM DTT. Fractions of interest were pooled, and 0.4 mg of TEV protease and 2 mM of fresh DTT were added to the sample, which was incubated 1 h at RT. Then, the sample was boiled 5 min at 95°C and spun down 5 min at 16,000 g at 4°C. It was concentrated using Novagen concentrators with 3.5 kDa cut off membranes centrifuged at 5,000 xg, and injected on a gel filtration column (Superdex 16/600 HiLoad 75 pg) equilibrated with DPBS (D1408; Sigma) at pH 7.2. Fractions were pooled, supplemented with 1 mM fresh DTT, and concentrated using a 3.5 kDa cut off concentrator centrifuged at 5,000 g. EDTA-free protease inhibitors (cOmplete EDTA-free, Sigma-Aldrich) were added at a final 1X concentration. The quality of the purified proteins was analyzed by SDS-PAGE and the protein concentrations were determined by spectrophotometry using the absorbance at 280 nm. All concentrated proteins were stored in small aliquots at −80 °C.

### Expression and purification of KIF2C and KIF2A N-terminal domains

The constructs 8His-GB1-TEVsite-KIF2C_1-79_ (WT and K52E/K54E) and 8His-GB1-TEVsite-KIF2A_1-77_KIF2C_1-79_ were expressed in *E. coli* BL21 (DE3) Star cells supplemented with 100 μg/ml ampicillin. Unlabeled proteins were produced by growing bacteria in 2x Luria broth (LB) media. ^15^N and/or ^13^C-labeled proteins were produced by growing the bacteria in M9 minimal medium containing 0.5 g/l ^15^NH4Cl and 2 g/l ^13^C-glucose. Expression was induced at an OD_400_ of 0.6-0.8 with 1 mM IPTG and cells were harvested after overnight of incubation at 20 °C by centrifugation. The bacterial pellet was resuspended in 35 mL of a solution containing 50 mM Tris-HCl at pH 8.0, 150 mM NaCl, 0.5% glycerol, 30 μL Triton X-100. Cells are sonicated on ice 2.5 min in total with 1s ON/9s OFF cycle of sonication (60% amplitude) at 10°C. 5 mM MgSO_4_, 0.5 μL benzonase (E1014; Millipore), 2 mM EDTA were added to the solution and incubated for 20 min at 4°C. Then, the lysate was clarified by centrifugation during 15 min at 15,000 g at 4°C. The soluble fraction was loaded on a Ni-NTA histidine-affinity column (5 mL HisTrap Excel, GE Healthcare) using a 1.5 mL/min flow rate. The column was washed with a solution containing 50 mM Tris HCl at pH 8.0, 150 mM NaCl, 5 mM EDTA. Then, the sample was eluted using an imidazole gradient over 45 mL, the final buffer containing 50 mM Tris HCl at pH 8.0, 150 mM NaCl, 1 M imidazole. The fractions of interest were pooled and dialyzed in a solution containing 50 mM Tris HCl at pH 8.0, 150 mM NaCl, 5 mM EDTA overnight. To cleave the tag, 0.4 mg of TEV protease, 1X of EDTA-free protease inhibitors (cOmplete EDTA-free) and 2 mM of fresh DTT were added to the sample, which was incubated 1 h at room temperature. The sample was then loaded on a HisTrap column and the tag-free proteins were collected in the flow through. The sample was concentrated using a Novagen concentrator with a 3.5 kDa cut off membrane centrifuged at 5,000 g, and injected on a gel filtration column (Superdex 16/600 HiLoad 200 pg) equilibrated with a buffer containing 50 mM Tris-HCl, 50 mM NaCl at pH 7.5 for non-labeled samples or DPBS (D1408; Sigma) at pH 7.0 for ^15^N and/or ^13^C-labeled samples. Fractions are pooled, concentrated using 3 kDa cut off concentrators centrifuged at 4,500 xg. The quality of the purified proteins was analyzed by SDS-PAGE 15% and the protein concentrations were determined by spectrophotometry using the absorbance at 280 nm. All concentrated proteins were stored in small aliquots at −80 °C.

### Search for BRCA2 partners by pull-down & mass spectrometry

Identification of phospho-dependent BRCA2 partners was carried out following our recently published protocol (Bouguechtouli *et al*., 2024). After production and purification of the 6His-Avi-BRCA2_167-260_ construct, half of the peptide sample was phosphorylated by adding PLK1 (provided by the recombinant protein platform of Institut Curie, Paris) at a molar ratio of about 200:1, as published (Alik et al., 2020; Julien et al., 2020a). Biotinylation of 6His-Avi-BRCA2_167-260_, either non-phosphorylated or phosphorylated, was performed by incubating the peptide at 100 µM in a solution containing 2 mM ATP, 600 µM biotin, 5 mM MgCl2, 1 mM DTT, 1X protease inhibitors, together with 0.7 µM of the BirA enzyme (produced and purified in the lab) in a buffer containing 50 mM HEPES at pH 7.0, 150 mM NaCl, 1 mM EDTA. The peptide was then incubated 1h 30 at RT and injected on a gel filtration column (Superdex 16/60 HiLoad 75 pg) previously equilibrated with a solution containing 50 mM HEPES pH 7.0, 75 mM NaCl, 1 mM EDTA. The fractions of interest were pooled, supplemented with 1 mM fresh DTT, and the sample was concentrated using a 3 kDa cut off concentrator centrifuged at 5,000 g. EDTA-free protease inhibitors (cOmplete EDTA-free; Sigma-Aldrich) were added at a final 1X concentration and the sample was flash frozen using liquid nitrogen.

From here, all conditions were performed with 5 replicates treated at the same time in order to favor the reproducibility of the experiment. Two ng of recombinant biotinylated 6His-Avi-BRCA2_167-260_ in Phosphate Buffered Saline (PBS) at pH 7.5, 1 mM DTT, 1X protease inhibitors were added to 50 µL of Streptavidin-coated magnetic beads (Streptavidin Mag-Beads; Genscript) in a final volume of 100 µL. Beads were incubated for 1 h at RT on a rotating wheel and were washed 3 times using 500 µL of PBS. They were then washed 2 times with 500 µL of a solution containing 50 mM HEPES at pH 7.2, 150 mM NaCl, 2 mM EDTA, 10 mM NaF, 0.5 mM PMSF, 1 X protease inhibitors, 1 mM DTT, 1X PhosphoSTOP mixed with 800 µg of lysed cells extracts (HEK293 cells synchronized or not with nocodazole) in 20 mM HEPES at pH 7.6, 150 mM NaCl, 0.1% NP40, 2 mM EGTA, 1.5 mM MgCl2, 50 mM NaF, 10% glycerol, 20 mM b-glycerophosphate, 1 mM DTT, 1X protease inhibitors. The beads were incubated for 2 h at RT on a rotating wheel. They were washed 3 times with 100 µL of a solution containing 50 mM HEPES at pH 7.2, 150 mM NaCl, 2 mM EDTA, 10 mM NaF, 0.5 mM PMSF, 1 mM DTT, and washed twice without resuspension of the beads with 500 µL of a solution containing 50 mM ammonium bicarbonate at pH 8.2. The beads were then kept on ice in 500 µL of the buffer containing 50 mM ammonium bicarbonate at pH 8.2, and analyzed by the mass spectrometry platform of Institut Curie (Paris). Further data processing was achieved on the website of the platform using the tool MYPROMS.

### NMR experiments

Most experiments were carried out at 10° C on 600 and 700 MHz Bruker spectrometers equipped with a triple resonance cryoprobe. Spectra were processed in Topspin 4.3.0 and analyzed with CcpNmr Analysis 2.4.2 software. NMR chemical shift assignments of the BRCA2 fragments used in this study are published (Julien *et al*., 2020b). To assign the NMR chemical shifts of the KIF2C and KIF2A N-terminal domains, standard 3D triple resonance NMR experiments (HNCO, HNCACO, CBCA(CO)HN, HNCA and HNCACB) were carried out on samples of proteins at 300 µM in DPBS pH 7.0, 2 mM DTT, 5% D_2_O, 50 mM DSS. Analysis of these experiments was performed, and the secondary structure of the KIF2C and KIF2A N-terminal domains was deduced from the resulting assigned backbone ^1^H, ^15^N and ^13^C chemical shifts using the website https://st-protein02.chem.au.dk/ncSPC/cgi-bin/selection_screen_ncSPC.py. To check the phosphorylation states of the ^15^N-labeled BRCA2 fragments, we carried out ^1^H-^15^N SOFAST-HMQC experiments on samples of proteins at 150 µM in DPBS pH 7.2, 2 mM DTT, 5% D_2_O, 50 mM DSS, mixed with 3 µM PLK1, 5 mM MgCl_2_, 2 mM ATP, 1X EDTA-free protease inhibitors.

To identify protein-protein interaction sites, we carried out ^15^N SOFAST-HMQC experiments (1536 x 200 timepoints, 256 scans, 50 ms of interscan delay) on 150 μL of ^15^N-labeled non-phosphorylated and phosphorylated BRCA2_167-270_ (and BRCA2_48-218_ in Figure 2C) mixed with non-labeled KIF2C_1-79_ and KIF2A_1-77_ at molar ratios of 1:0, 1:1, in DPBS pH 7.2. In addition, we performed ^15^N SOFAST-HMQC experiments (1536 x 160 timepoints, 200 scans, 40 ms of interscan delay) on 150 μl of ^15^N-labeled KIF2C_1-79_ (WT or K52E/K54E) and KIF2A_1-77_ mixed with non-phosphorylated and phosphorylated BRCA2 peptides (pT207) and Sgo2 (pT537, pT620) at molar ratios of 1:0, 1:0.5, 1:1, 1:2, in DPBS pH 7.0. The peptides used for all experiments were synthesized by GenScript (Piscataway, NY) (Suppl. Table 3).

To measure the chemical shifts of the BRCA2 peptide bound state, we recorded ^15^N D-CEST experiments at 10°C on a 800 MHz Bruker spectrometer (Yuwen et al., 2018). The NMR sample consisted of ^15^N-labeled BRCA2_167-260_ and BRCA2_48-218_ at 200 µM in the presence of unlabeled KIF2C_1-79_ at a molar ratio 10:1, in DPBS pH 7.2. We used effective DANTE B1 fields of 10, 20 and 40 Hz with the position of the RF field varied over 500, 800 and 1500 Hz (*sw*_CEST_), respectively, using step sizes of 20.8, 36.4 and 65.2 Hz. The D-CEST period, T_Ex_, was set to 300 ms; each 2D plane was recorded with 16 transients per FID. The prescan delay was 1.5 s, and (795,115) complex points in (t2, t1) were recorded, to give a net acquisition time of ∼2 h per spectrum. Total measurement time for D-CEST experiments was ∼2 d per B1 field, so altogether ∼6 d. All NMR exchange data were fitted simultaneously using the program ChemEx.

### Isothermal titration calorimetry (ITC)

ITC measurements were performed with the KIF2C_1-79_ or KIF2A_1-77_ protein at 8-10 µM in the cell and the BRCA2 or Sgo2 peptide (Suppl. Table 3) at 80-100 µM in the syringe. The buffer was 50 mM Tris-HCl, pH 7.5, 50 mM NaCl. The experiments were carried out on a VP-ITC instrument (Malvern), using automatic injections of 8 or 10 µl at 20°C. The peptides used for all experiments were synthesized by GenScript (Piscataway, NY) (Suppl. Table 3). Control experiments were carried out by injecting peptides into the cell filled with buffer, to estimate the heat of dilution. The titration curves were analyzed using the program Origin 7.0 (OriginLab) and fitted to a one-site binding model.

### Size-Exclusion Chromatography coupled to multi-angle light scattering (SEC-MALS)

SEC-MALS was used to measure the molecular mass of the N-terminal domain of KIF2C in solution. KIF2C N-terminal domain was loaded on an Agilent Technologies HPLC system with a BIOSEC 3-300 column (4,6×300mm) (flow rate at 200 μl/min). The light scattering was measured with a 3 angles MALS system from WYATT company and the RI (Refractive Index) to analyze the difference of refraction index is a VISCOTEK (MALVERN) equipment. The chromatography buffer was 25 mM Tris-HCl buffer (pH 7.0), 50 mM NaCl. A calibration was performed with BSA as a standard. Data were analyzed using the ASTRA software. To represent the data, the normalized absorbance at 280 nm was overlaid with the molar mass (Da), and both parameters were plotted as a function of the elution volume.

### AlphaFold calculations

AlphaFold models of full-length monomeric proteins were obtained from the AlphaFold Protein Structure Database (Jumper et al., 2021; Varadi et al., 2022). Models of protein-peptide complexes were computed using the server of the Integrative Bioinformatics platform of I2BC (https://bioi2.i2bc.paris-saclay.fr). For each complex, a series of 15 models were calculated and the models were analyzed using the delivered heatmap and lDDT plots, as well as the pTM and ipTM scores (Evans et al., 2022; Jumper *et al*., 2021).

### Cell extracts, GFP-TRAP, and western blotting for identifying BRCA2-KIF2C interaction in cells

For analysis of the complex between BRCA2 and KIF2C in mitosis, DLD1 BRCA2-/- cells stably expressing EGFP-MBP-BRCA2 WT were synchronized with 0.1 μg/ml nocodazole (Sigma Aldrich) for 15 h and 30 min. The mitotic population was enriched by shake off. Cell pellets were harvested and lysed with extraction buffer A (20 mM HEPES (pH 7.5), 150 mM NaCl, 0.1% NP_4_0, 2 mM EGTA, 1.5 mM MgCl2, 50 mM NaF, 10% glycerol, 1 mM Na_3_VO_4_, 20 mM ß-glycerophosphate, 1 mM DTT and EDTA-free Protease Inhibitor Cocktail). Samples were pre-cleared by centrifugation at maximum speed for 30 min. GFP-TRAP agarose beads (ChromoTek) were equilibrated with extraction buffer A before being incubated with mitotic lysates for 1.5 h at 4°C. The beads were then washed 4 times with extraction buffer A and 3 times with extraction buffer A containing 250 mM NaCl. Then, beads were heated at 95°C for 5 min in 3X SDS-PAGE Laemmli buffer and spun again. The supernatant containing the eluted proteins were then migrated on a 4-15% gradient pre-cast SDS-PAGE (Bio-Rad), transferred onto nitrocellulose membranes (Amersham) and finally blotted with the following antibodies: anti-mouse BRCA2 (1:1000, OP95) and anti-mouse KIF2C (1:1000).

### Metaphase spreads coupled with immunofluorescence

To study the localization of BRCA2- pT207 and KIF2C at the kinetochores, cells stably expressing BRCA2 WT were synchronized using 0.1μg/ml nocodazole for 4 h to enrich the mitotic population. Dividing cells were collected by shake off, pelleted, and subjected to a hypotonic shock with 50 mM KCl for 20 min. Cells were then spun onto Poly-D-lysine (Sigma Aldrich) precoated coverslips for 5 min at 500 g. Then cells were fixed and permeabilized simultaneously with 4% PFA, 0.5% Triton, 50mM NaF, 20mM ß -glycerophosphate and 1mM Na_3_VO_4_ for 20 min at RT. After 3 washes of PBS-Tween 0.2%, 5 min each, coverslips were blocked with PBS-BSA 4% for 30 min at RT before being incubated with the following antibodies overnight at 4°C: rabbit anti-pBRCA2-T207 (as previously described (Ehlen *et al*., 2020) (1:500, Genscript), mouse anti-KIF2C (1:100) and human anti-Crest (1:100). Coverslips were washed twice for 5 min each with PBS-Triton 0.1% and incubated for 2h at RT with the appropriate Alexa Fluor secondary antibody: donkey anti-rabbit Alexa-488 (1:1000), donkey anti-mouse Alexa-488 (1:1000), donkey anti-rabbit Alexa-594 (1:1000) or goat anti-human Alexa-555 (1:1000). Primary and secondary antibodies were diluted with PBS-Tween 0.1%, 5% BSA. After 2 washes of the secondary antibodies with PBS-Triton 0.1% and a final wash with PBS, coverslips were stained with DAPI (1μg/ml, Merck, Cat. # 268298) and mounted using Prolong Glass antifade (P-36984, Thermo Fisher Scientific). Images were acquired with a Laser Scanning Confocal Microscope LSM900 coupled to an upright Axio Imager 2 Microscope (Zeiss). Image acquisition was done with a 100x oil objective.

### Optogenetic activation of KIF2C condensation

Cells were plated at around 70% confluency in DMEM. Expression of Opto-KIF2C was induced with 1 μg/ml doxycycline for 16 h. In the case of studying cells in mitosis, 100 ng/ml nocodazole was added for 16 h. To activate the light, the samples were transferred into a custom-made illumination box containing an array of 24 LEDs (488 nm) delivering 10 mW/cm^2^. Cry2 oligomerization was induced using 5 min of light-dark cycles (4 s light followed by 10 s dark).

### Pulldown of biotinylated proteins

TurboID Flp-In 293 T-REx cell lines stably transfected with opto-KIF2C recombinant protein and grown to 75% confluence were incubated with 6 ng/ml of doxycycline for 16 hours. The next day, the cells were incubated with 500 mM of biotin for 15 min. Cells were then washed with PBS and lysed with lysis buffer (50 mM Tris-HCl pH 7.5, 150 mM NaCl, 1 mM EDTA, 1 mM EGTA, 1% NP-40, 0.2% SDS, 0.5% sodium deoxycholate) supplemented with 1X complete protease inhibitor, 1X phosphatase inhibitor and 250U benzonase. Lysed cells were incubated on a rotating wheel for 1 hour at 4°C prior sonication on ice (40% amplitude, 3 cycles 10 sec sonication-2 sec resting). After centrifugation (7750 rcf.) for 30 min at 4°C, the cleared supernatant was transferred to a new tube and the total protein concentration was determined using the Bradford protein assay. For each condition, 1 mg of proteins was incubated with 50 ml of streptavidin-Agarose beads on a rotating wheel at 4°C for 3 hours. After 1 min centrifugation (400 rcf.), the beads were washed sequentially with 1 ml of lysis buffer, 1 ml wash buffer 1 (2% SDS in H_2_O), 1 ml wash buffer 2 (0.2% sodium deoxycholate, 1% Triton X-100, 500 mM NaCl, 1 mM EDTA, and 50 mM HEPES pH 7.5), 1 ml Wash Buffer 3 (250 mM LiCl, 0.5% NP-40, 0.5% sodium deoxycholate, 1 mM EDTA, 500 mM NaCl and 10 mM Tris pH 8) and 1 ml Wash Buffer 4 (50 mM Tris pH 7.5 and 50 mM NaCl). For Western blot analysis of KIF2C partners enriched in optogenetic KIF2C condensates, cells were simultaneously incubated with 500 mM of biotin and exposed to blue light for 15 min of light-dark cycles (4 s light followed by 30 s dark). Biotin proximity labeling of light induced KIF2C partners was performed using streptavidin-coated beads as described previously. Bound proteins were eluted from the agarose beads with 80 ml of 2X Laemmli sample buffer and incubated at 95°C for 10 min. 5mg of the lysates were used for Western blot analysis and probed by immunoblotting to detect proteins that are associated with KIF2C clusters.

### Western blotting for TurboID constructs

Constructs used for western blotting are listed in Suppl. Table S2. Whole cell extracts were prepared by lysing cells in RIPA buffer (50 mM Tris-HCl at pH 8.0, 150 mM NaCl, 1% NP-40, 1% deoxycholate, 0.1% SDS) for 30 min on ice. After centrifugation, the supernatant was collected and analyzed for protein amount using the Quick Start Bradford protein assay kit. After addition of Laemmli Sample buffer 1X (Biorad, C161-0737), the sample was heated for 5 minutes at 95°C. According to the manufacturer’s instructions (BioRad), 40 μg of protein samples were resolved on precast SDS-PAGE gels (4-15% and 10%) and transferred to a nitrocellulose membrane using the Bio-Rad Trans-Blot Turbo transfer device. Membranes were saturated with 10% non-fat milk diluted in TBS-0.2% Tween 20 (TBS-T), incubated with primary antibodies (Suppl. Table 4) overnight at 4°C and with secondary anti-mouse-HRP (1 :1000) or anti-rabbit-HRP (1:1000) antibodies for 1h. Blots were developed with ECL according to the manufacturer’s instructions.

### Immunofluorescence staining and Fluorescent Microscopy Imaging

Cells grown on coverslips were fixed with PBS/4% paraformaldehyde (PFA) for 15 min at RT followed by a 10 min permeabilization step in PBS/0.2% Triton X-100-PBS and blocked in PBS/3% BSA for 30 min. An intermediate wash with PBS 1x for 5 min was used to completely remove buffers. For immunofluorescence staining, primary antibodies (Suppl. Table 4) were diluted in blocking solution and incubated for 1 h at RT, after which cells were washed using 1X PBS. Next, the corresponding secondary antibodies coupled to the fluorochrome were diluted in a blocking solution and also incubated for 1 h at RT. DNA was stained with Hoechst 33258 (Invitrogen, Cat H21491) during 5 min at RT. Coverslips were then mounted onto glass slides using Prolong Gold Antifade Reagent (Invitrogen, Cat P36930). The finished coverslips were stored at 4°C. Images were acquired on a LEICA SP8X inverted confocal laser scanning microscope equipped with a 63x HC Plan Apochromat CS2 oil-immersion objective (NA: 1.4) (Leica) with hybrid GaAsP detectors (Hamamatsu). A white light laser was used at 405, 488 and 561 nm wavelengths to excite DAPI, GFP and mCherry respectively and their fluorescence was collected through the following bandpass 415-461, 495-550 and 575-800 nm.

### FRAP experiments

For FRAP experiments, we used a Leica TCS SP8X system equipped with a 63x HC Plan Apochromat CS2 oil-immersion objective (NA: 1.4) (Leica), 488 nm. Cells were preseeded in a μ-Slide 8 Well high Glass Botton 170 μm (Ibidi, 80807). They were incubated for 12 h in the presence of 1 μg/ml doxycycline to induce opto-KIF2C expression and nocodazole was added to block cells in mitosis at 37°C. The whole living cells were illuminated to primary activate foci for one cycle of light (15 s followed by dark for 30 s), and then a plane about 0.5 μm thick showing a large number of foci was exposed during 20 min with light-dark cycles (8 s light followed by 30 s dark). To maintain cell viability, all experiments were carried out at 37 °C and 5 % CO_2_. FRAP acquisitions were taken using 3D imaging and a 3 Airy pinhole, because the condensates move very fast and we did not want to lose them or to confuse the recovery with a movement on itself. The bleaching events were performed in a circular region with a diameter of 500 nm and monitored over time for fluorescence recovery. In order not to bleach the entire condensate, we targeted its boundaries. The laser was set at 2 % for bleaching and 20 % for imaging with a scanning speed of 400 Hz. The FRAP sequence was composed of a short pre-bleach sequence of 3 images, the bleaching event during 2.61 s, and 1 post-bleach sequence repeated 100-300 times. To collect the images, we used a PMT with a gain fixed at 400 and a bandpass 576-800 (not to lose fluorescence). Since we only had one marking and no auto-fluorescence, FRAP curves were independently corrected and processed to obtain a double normalization as follow: the mean intensity of every bleached region was measured and the background intensity was subtracted by measuring a region outside the cell. Acquisition-related bleaching correction was performed by dividing values by the whole cell. Then, to display the recovery curves from 0 to 1, normalization was performed using the average of the pre-bleached signal and the 1st post-bleached value. As the recovery curves display a biphasic aspect, the mean curve was fitted by a double exponential function to extract both half-time recovery 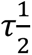 and the mobile fraction of pooled recovery curves.

### Structured illumination microscopy (SIM)

SIM experiments were performed on a ZEISS Elyra 7 – Lattice SIM equipped with a 63x Plan-Apochromat (N.A. 1.4) oil immersion objective and coupled with a PCO.edge sCMOS camera (pixel size: 6.5 µm; bit depth 16 bit). For Alexa 568 imaging, excitation was performed with a 561 nm laser (100 mW), and fluorescence collected using a dual band emission filter (BP 495-550 nm, BP 570-620 nm). For Alexa 488 imaging, excitation was performed with a 488 nm laser (100 mW, OPSL), and fluorescence collected using a triple band emission filter (BP 420-480 nm, BP 495-525 nm, LP 650 nm). For DAPI imaging, excitation was performed with a 405 nm laser (50 mW), and fluorescence collected using a triple band emission filter (BP 420-480 nm, BP 495-525 nm, LP 650 nm). For all the channels the illumination was structured as a lattice pattern (G4, 32 µm) and 13 phases were shifted for each plane and a z-step of 273 nm was used to generate 3D-SIM acquisitions (leap mode). SIM processing was performed on ZEN Black (ZEISS, version 16.0) and to correct chromatic aberrations, alignment procedure (ZEN Black) was applied on both channels after measurements on multispectral calibration beads.

### Image and statistical analysis

Image treatment and analysis were performed using Imaris version 10.1.0 and FIJI version 2.0.0 software (Schindelin et al., 2012). The plugin ezcolocalization for FIJI software was used to evaluate colocalization in 2D and 3D (Stauffer et al., 2018). To quantify colocalization, Pearson Correlation Coefficients were calculated. DAPI was used as a mask in all experiments except for colocalization between KIF2C and BRCA2 pT207. In the case of colocalization between KIF2C and pBRCA2 pT207, a mask based on KIF2C condensates was used. The analysis of the FRAP experiment was carried out using a plugin for FIJI developed at I2BC. In all graphs, error bars represent the standard deviation (SD) from 2 or 3 independent experiments, and scatter dot plots show median with 95% CI. Statistical significance of differences was calculated with unpaired two-tailed t-test of Mann–Whitney by using GraphPad Prism version 10.1.1. Graphs were generated using GraphPad Prism or SuperPlotsOfData by Huygens.

### Time-lapse video microscopy

To observe the fusion events, cells expressing KIF2C WT (+D+L conditions) were illuminated under the microscope using a protocol similar to that described for FRAP. Images were collected in 3-5 z-stacks with a step of 0.5 μm every 5 s. Further 3D visualization and measurement of the volume of the condensates were performed using Imaris software version 10.1.0 with a PSF correction of 0.6785.

## References

Alghoul, E., Basbous, J., and Constantinou, A. (2021). An optogenetic proximity labeling approach to probe the composition of inducible biomolecular condensates in cultured cells. STAR Protoc 2, 100677. 10.1016/j.xpro.2021.100677.

Alik, A., Bouguechtouli, C., Julien, M., Bermel, W., Ghouil, R., Zinn-Justin, S., and Theillet, F.X. (2020). Sensitivity-Enhanced (13) C-NMR Spectroscopy for Monitoring Multisite Phosphorylation at Physiological Temperature and pH. Angew Chem Int Ed Engl 59, 10411–10415. 10.1002/anie.202002288.

Andrews, P.D., Ovechkina, Y., Morrice, N., Wagenbach, M., Duncan, K., Wordeman, L., and Swedlow, J.R. (2004). Aurora B regulates MCAK at the mitotic centromere. Dev Cell 6, 253–268. 10.1016/s1534-5807(04)00025-5.

Bakhoum, S.F., Genovese, G., and Compton, D.A. (2009a). Deviant kinetochore microtubule dynamics underlie chromosomal instability. Curr Biol 19, 1937–1942. 10.1016/j.cub.2009.09.055.

Bakhoum, S.F., Thompson, S.L., Manning, A.L., and Compton, D.A. (2009b). Genome stability is ensured by temporal control of kinetochore-microtubule dynamics. Nat Cell Biol 11, 27–35. 10.1038/ncb1809.

Bodmer, B.S., Breithaupt, A., Heung, M., Brunetti, J.E., Henkel, C., Muller-Guhl, J., Rodriguez, E., Wendt, L., Winter, S.L., Vallbracht, M., et al. (2023). In vivo characterization of the novel ebolavirus Bombali virus suggests a low pathogenic potential for humans. Emerg Microbes Infect 12, 2164216. 10.1080/22221751.2022.2164216.

Bouguechtouli, C., Ghouil, R., Alik, A., Dingli, F., Loew, D., and Theillet, F.X. (2024). Structural characterization of stem cell factors Oct4, Sox2, Nanog and Esrrb disordered domains, and a method to identify their phospho-dependent binding partners. Comptes Rendus – Chimie Acad Sci.

Branon, T.C., Bosch, J.A., Sanchez, A.D., Udeshi, N.D., Svinkina, T., Carr, S.A., Feldman, J.L., Perrimon, N., and Ting, A.Y. (2018). Efficient proximity labeling in living cells and organisms with TurboID. Nat Biotechnol 36, 880–887. 10.1038/nbt.4201.

Bugaj, L.J., Choksi, A.T., Mesuda, C.K., Kane, R.S., and Schaffer, D.V. (2013). Optogenetic protein clustering and signaling activation in mammalian cells. Nat Methods 10, 249–252. 10.1038/nmeth.2360.

Carmena, M., Pinson, X., Platani, M., Salloum, Z., Xu, Z., Clark, A., Macisaac, F., Ogawa, H., Eggert, U., Glover, D.M., et al. (2012). The chromosomal passenger complex activates Polo kinase at centromeres. PLoS Biol 10, e1001250. 10.1371/journal.pbio.1001250.

Cimini, D., Howell, B., Maddox, P., Khodjakov, A., Degrassi, F., and Salmon, E.D. (2001). Merotelic kinetochore orientation is a major mechanism of aneuploidy in mitotic mammalian tissue cells. J Cell Biol 153, 517–527. 10.1083/jcb.153.3.517.

Corno, A., Cordeiro, M.H., Allan, L.A., Lim, Q.W., Harrington, E., Smith, R.J., and Saurin, A.T. (2023). A bifunctional kinase-phosphatase module balances mitotic checkpoint strength and kinetochore-microtubule attachment stability. EMBO J 42, e112630. 10.15252/embj.2022112630.

Ehlen, A., Martin, C., Miron, S., Julien, M., Theillet, F.X., Ropars, V., Sessa, G., Beaurepere, R., Boucherit, V., Duchambon, P., et al. (2020). Proper chromosome alignment depends on BRCA2 phosphorylation by PLK1. Nat Commun 11, 1819. 10.1038/s41467-020-15689-9.

Elowe, S., Hummer, S., Uldschmid, A., Li, X., and Nigg, E.A. (2007). Tension-sensitive Plk1 phosphorylation on BubR1 regulates the stability of kinetochore microtubule interactions. Genes Dev 21, 2205–2219. 10.1101/gad.436007.

Evans, R., O’Neill, M., Pritzel, A., Antropova, N., Senior, A., Green, T., Žídek, A., Bates, R., Blackwell, S., Yim, J., et al. (2022). Protein complex prediction with AlphaFold-Multimer. bioRxiv, 2021.2010.2004.463034. 10.1101/2021.10.04.463034.

Fairhead, M., and Howarth, M. (2015). Site-specific biotinylation of purified proteins using BirA. Methods Mol Biol 1266, 171–184. 10.1007/978-1-4939-2272-7_12.

Frattini, C., Promonet, A., Alghoul, E., Vidal-Eychenie, S., Lamarque, M., Blanchard, M.P., Urbach, S., Basbous, J., and Constantinou, A. (2021). TopBP1 assembles nuclear condensates to switch on ATR signaling. Mol Cell 81, 1231–1245 e1238. 10.1016/j.molcel.2020.12.049.

Gayatri, S., and Bedford, M.T. (2014). Readers of histone methylarginine marks. Biochim Biophys Acta 1839, 702–710. 10.1016/j.bbagrm.2014.02.015.

Godek, K.M., Kabeche, L., and Compton, D.A. (2015). Regulation of kinetochore-microtubule attachments through homeostatic control during mitosis. Nat Rev Mol Cell Biol 16, 57–64. 10.1038/nrm3916.

Hernandez-Vega, A., Braun, M., Scharrel, L., Jahnel, M., Wegmann, S., Hyman, B.T., Alberti, S., Diez, S., and Hyman, A.A. (2017). Local Nucleation of Microtubule Bundles through Tubulin Concentration into a Condensed Tau Phase. Cell Rep 20, 2304–2312. 10.1016/j.celrep.2017.08.042.

Hunter, A.W., Caplow, M., Coy, D.L., Hancock, W.O., Diez, S., Wordeman, L., and Howard, J. (2003). The kinesin-related protein MCAK is a microtubule depolymerase that forms an ATP-hydrolyzing complex at microtubule ends. Mol Cell 11, 445–457. 10.1016/s1097-2765(03)00049-2.

Jiang, H., Wang, S., Huang, Y., He, X., Cui, H., Zhu, X., and Zheng, Y. (2015). Phase transition of spindle-associated protein regulate spindle apparatus assembly. Cell 163, 108–122. 10.1016/j.cell.2015.08.010.

Jiang, X., Ho, D.B.T., Mahe, K., Mia, J., Sepulveda, G., Antkowiak, M., Jiang, L., Yamada, S., and Jao, L.E. (2021). Condensation of pericentrin proteins in human cells illuminates phase separation in centrosome assembly. J Cell Sci 134. 10.1242/jcs.258897.

Joukov, V., and De Nicolo, A. (2018). Aurora-PLK1 cascades as key signaling modules in the regulation of mitosis. Sci Signal 11. 10.1126/scisignal.aar4195.

Julien, M., Bouguechtouli, C., Alik, A., Ghouil, R., Zinn-Justin, S., and Theillet, F.X. (2020a). Multiple Site-Specific Phosphorylation of IDPs Monitored by NMR. Methods Mol Biol 2141, 793–817. 10.1007/978-1-0716-0524-0_41.

Julien, M., Ghouil, R., Petitalot, A., Caputo, S.M., Carreira, A., and Zinn-Justin, S. (2021). Intrinsic Disorder and Phosphorylation in BRCA2 Facilitate Tight Regulation of Multiple Conserved Binding Events. Biomolecules 11. 10.3390/biom11071060.

Julien, M., Miron, S., Carreira, A., Theillet, F.X., and Zinn-Justin, S. (2020b). (1)H, (13)C and (15)N backbone resonance assignment of the human BRCA2 N-terminal region. Biomol NMR Assign 14, 79–85. 10.1007/s12104-019-09924-8.

Jumper, J., Evans, R., Pritzel, A., Green, T., Figurnov, M., Ronneberger, O., Tunyasuvunakool, K., Bates, R., Zidek, A., Potapenko, A., et al. (2021). Highly accurate protein structure prediction with AlphaFold. Nature 596, 583–589. 10.1038/s41586-021-03819-2.

King, M.R., and Petry, S. (2020). Phase separation of TPX2 enhances and spatially coordinates microtubule nucleation. Nat Commun 11, 270. 10.1038/s41467-019-14087-0.

Kleyman, M., Kabeche, L., and Compton, D.A. (2014). STAG2 promotes error correction in mitosis by regulating kinetochore-microtubule attachments. J Cell Sci 127, 4225–4233. 10.1242/jcs.151613.

Kruse, T., Zhang, G., Larsen, M.S., Lischetti, T., Streicher, W., Kragh Nielsen, T., Bjorn, S.P., and Nilsson, J. (2013). Direct binding between BubR1 and B56-PP2A phosphatase complexes regulate mitotic progression. J Cell Sci 126, 1086–1092. 10.1242/jcs.122481.

Lan, W., Zhang, X., Kline-Smith, S.L., Rosasco, S.E., Barrett-Wilt, G.A., Shabanowitz, J., Hunt, D.F., Walczak, C.E., and Stukenberg, P.T. (2004). Aurora B phosphorylates centromeric MCAK and regulates its localization and microtubule depolymerization activity. Curr Biol 14, 273–286. 10.1016/j.cub.2004.01.055.

Lawrence, E.J., Chatterjee, S., and Zanic, M. (2023). More is different: Reconstituting complexity in microtubule regulation. J Biol Chem 299, 105398. 10.1016/j.jbc.2023.105398.

Lee, M., Daniels, M.J., and Venkitaraman, A.R. (2004). Phosphorylation of BRCA2 by the Polo-like kinase Plk1 is regulated by DNA damage and mitotic progression. Oncogene 23, 865–872. 10.1038/sj.onc.1207223.

Lenart, P., Petronczki, M., Steegmaier, M., Di Fiore, B., Lipp, J.J., Hoffmann, M., Rettig, W.J., Kraut, N., and Peters, J.M. (2007). The small-molecule inhibitor BI 2536 reveals novel insights into mitotic roles of polo-like kinase 1. Curr Biol 17, 304–315. 10.1016/j.cub.2006.12.046.

Li, H., Liu, X.S., Yang, X., Wang, Y., Wang, Y., Turner, J.R., and Liu, X. (2010). Phosphorylation of CLIP-170 by Plk1 and CK2 promotes timely formation of kinetochore-microtubule attachments. EMBO J 29, 2953–2965. 10.1038/emboj.2010.174.

Lin, H.R., Ting, N.S., Qin, J., and Lee, W.H. (2003). M phase-specific phosphorylation of BRCA2 by Polo-like kinase 1 correlates with the dissociation of the BRCA2-P/CAF complex. J Biol Chem 278, 35979–35987. 10.1074/jbc.M210659200.

Maney, T., Hunter, A.W., Wagenbach, M., and Wordeman, L. (1998). Mitotic centromere-associated kinesin is important for anaphase chromosome segregation. J Cell Biol 142, 787–801. 10.1083/jcb.142.3.787.

Maney, T., Wagenbach, M., and Wordeman, L. (2001). Molecular dissection of the microtubule depolymerizing activity of mitotic centromere-associated kinesin. J Biol Chem 276, 34753–34758. 10.1074/jbc.M106626200.

Maurer-Stroh, S., Dickens, N.J., Hughes-Davies, L., Kouzarides, T., Eisenhaber, F., and Ponting, C.P. (2003). The Tudor domain ‘Royal Family’: Tudor, plant Agenet, Chromo, PWWP and MBT domains. Trends Biochem Sci 28, 69–74. 10.1016/S0968-0004(03)00004-5.

Miesch, J., Wimbish, R.T., Velluz, M.C., and Aumeier, C. (2023). Phase separation of +TIP networks regulates microtubule dynamics. Proc Natl Acad Sci U S A 120, e2301457120. 10.1073/pnas.2301457120.

Moore, A.T., Rankin, K.E., von Dassow, G., Peris, L., Wagenbach, M., Ovechkina, Y., Andrieux, A., Job, D., and Wordeman, L. (2005). MCAK associates with the tips of polymerizing microtubules. J Cell Biol 169, 391–397. 10.1083/jcb.200411089.

Ogawa, T., Nitta, R., Okada, Y., and Hirokawa, N. (2004). A common mechanism for microtubule destabilizers-M type kinesins stabilize curling of the protofilament using the class-specific neck and loops. Cell 116, 591–602. 10.1016/s0092-8674(04)00129-1.

Ritter, A., Kreis, N.N., Louwen, F., Wordeman, L., and Yuan, J. (2015). Molecular insight into the regulation and function of MCAK. Crit Rev Biochem Mol Biol 51, 228–245. 10.1080/10409238.2016.1178705.

Sanhaji, M., Friel, C.T., Kreis, N.N., Kramer, A., Martin, C., Howard, J., Strebhardt, K., and Yuan, J. (2010). Functional and spatial regulation of mitotic centromere-associated kinesin by cyclin-dependent kinase 1. Mol Cell Biol 30, 2594–2607. 10.1128/MCB.00098-10.

Sanhaji, M., Ritter, A., Belsham, H.R., Friel, C.T., Roth, S., Louwen, F., and Yuan, J. (2014). Polo-like kinase 1 regulates the stability of the mitotic centromere-associated kinesin in mitosis. Oncotarget 5, 3130–3144. 10.18632/oncotarget.1861.

Saurin, A.T. (2018). Kinase and Phosphatase Cross-Talk at the Kinetochore. Front Cell Dev Biol 6, 62. 10.3389/fcell.2018.00062.

Schindelin, J., Arganda-Carreras, I., Frise, E., Kaynig, V., Longair, M., Pietzsch, T., Preibisch, S., Rueden, C., Saalfeld, S., Schmid, B., et al. (2012). Fiji: an open-source platform for biological-image analysis. Nat Methods 9, 676–682. 10.1038/nmeth.2019.

Selenko, P., Sprangers, R., Stier, G., Buhler, D., Fischer, U., and Sattler, M. (2001). SMN tudor domain structure and its interaction with the Sm proteins. Nat Struct Biol 8, 27–31. 10.1038/83014.

Shao, H., Huang, Y., Zhang, L., Yuan, K., Chu, Y., Dou, Z., Jin, C., Garcia-Barrio, M., Liu, X., and Yao, X. (2015). Spatiotemporal dynamics of Aurora B-PLK1-MCAK signaling axis orchestrates kinetochore bi-orientation and faithful chromosome segregation. Sci Rep 5, 12204. 10.1038/srep12204.

Siegel, J.J., and Amon, A. (2012). New insights into the troubles of aneuploidy. Annu Rev Cell Dev Biol 28, 189–214. 10.1146/annurev-cellbio-101011-155807.

Stauffer, W., Sheng, H., and Lim, H.N. (2018). EzColocalization: An ImageJ plugin for visualizing and measuring colocalization in cells and organisms. Sci Rep 8, 15764. 10.1038/s41598-018-33592-8.

Suijkerbuijk, S.J., Vleugel, M., Teixeira, A., and Kops, G.J. (2012). Integration of kinase and phosphatase activities by BUBR1 ensures formation of stable kinetochore-microtubule attachments. Dev Cell 23, 745–755. 10.1016/j.devcel.2012.09.005.

Talapatra, S.K., Harker, B., and Welburn, J.P. (2015). The C-terminal region of the motor protein MCAK controls its structure and activity through a conformational switch. Elife 4. 10.7554/eLife.06421.

Tanno, Y., Kitajima, T.S., Honda, T., Ando, Y., Ishiguro, K., and Watanabe, Y. (2010). Phosphorylation of mammalian Sgo2 by Aurora B recruits PP2A and MCAK to centromeres. Genes Dev 24, 2169–2179. 10.1101/gad.1945310.

Varadi, M., Anyango, S., Deshpande, M., Nair, S., Natassia, C., Yordanova, G., Yuan, D., Stroe, O., Wood, G., Laydon, A., et al. (2022). AlphaFold Protein Structure Database: massively expanding the structural coverage of protein-sequence space with high-accuracy models. Nucleic Acids Res 50, D439–D444. 10.1093/nar/gkab1061.

Wang, W., Cantos-Fernandes, S., Lv, Y., Kuerban, H., Ahmad, S., Wang, C., and Gigant, B. (2017). Insight into microtubule disassembly by kinesin-13s from the structure of Kif2C bound to tubulin. Nat Commun 8, 70. 10.1038/s41467-017-00091-9.

Weaver, B.A., and Cleveland, D.W. (2007). Aneuploidy: instigator and inhibitor of tumorigenesis. Cancer Res 67, 10103–10105. 10.1158/0008-5472.CAN-07-2266.

Wimbish, R.T., and DeLuca, J.G. (2020). Hec1/Ndc80 Tail Domain Function at the Kinetochore-Microtubule Interface. Front Cell Dev Biol 8, 43. 10.3389/fcell.2020.00043.

Woodruff, J.B., Ferreira Gomes, B., Widlund, P.O., Mahamid, J., Honigmann, A., and Hyman, A.A. (2017). The Centrosome Is a Selective Condensate that Nucleates Microtubules by Concentrating Tubulin. Cell 169, 1066–1077 e1010. 10.1016/j.cell.2017.05.028.

Wordeman, L., and Mitchison, T.J. (1995). Identification and partial characterization of mitotic centromere-associated kinesin, a kinesin-related protein that associates with centromeres during mitosis. J Cell Biol 128, 95–104. 10.1083/jcb.128.1.95.

Wu, Y.O., Bryant, A.T., Nelson, N.T., Madey, A.G., Fernandes, G.F., and Goodson, H.V. (2021). Overexpression of the microtubule-binding protein CLIP-170 induces a +TIP network superstructure consistent with a biomolecular condensate. PLoS One 16, e0260401. 10.1371/journal.pone.0260401.

Yuwen, T., Kay, L.E., and Bouvignies, G. (2018). Dramatic Decrease in CEST Measurement Times Using Multi-Site Excitation. Chemphyschem 19, 1707–1710. 10.1002/cphc.201800249.

Zhang, L., Shao, H., Huang, Y., Yan, F., Chu, Y., Hou, H., Zhu, M., Fu, C., Aikhionbare, F., Fang, G., et al. (2011). PLK1 phosphorylates mitotic centromere-associated kinesin and promotes its depolymerase activity. J Biol Chem 286, 3033–3046. 10.1074/jbc.M110.165340.

Zhang, X., Ems-McClung, S.C., and Walczak, C.E. (2008). Aurora A phosphorylates MCAK to control ran-dependent spindle bipolarity. Mol Biol Cell 19, 2752–2765. 10.1091/mbc.e08-02-0198.

